# Chemotherapy-Induced Cachexia and Model-Informed Dosing to Preserve Lean Mass in Cancer Treatment

**DOI:** 10.1101/2021.10.01.462698

**Authors:** Suzan Farhang-Sardroodi, Michael A. La Croix, Kathleen P. Wilkie

## Abstract

Although chemotherapy is a standard treatment for cancer, it comes with significant side effects. In particular, certain agents can induce severe muscle loss, known as cachexia, worsening patient quality of life and treatment outcomes. 5-fluorouracil, an anti-cancer agent used to treat several cancers, has been shown to cause muscle loss. Experimental data indicates a non-linear dose-dependence for muscle loss in mice treated with daily or week-day schedules. We present a mathematical model of chemotherapy-induced muscle wasting that captures this non-linear dose-dependence. Area-under-the-curve metrics are proposed to quantify the treatment’s effects on lean mass and tumour control. Model simulations are used to explore alternate dosing schedules, aging effects, and morphine use in chemotherapy treatment with the aim of better protecting lean mass while actively targeting the tumour, ultimately leading to improved personalization of treatment planning and improved patient quality of life.

**Author Summary:** In this paper we present a novel mathematical model for muscle loss due to cancer chemotherapy treatment. Loss of muscle mass relates to increased drug toxicity and side-effects, and to decreased patient quality of life and survival rates. With our model, we examine the therapeutic efficacy of various dosing schedules with the aim of controlling a growing tumour while also preserving lean mass. Preservation of body composition, in addition to consideration of inflammation and immune interactions, the gut microbiome, and other systemic health measures, may lead to improved patient-specific treatment plans that improve patient quality of life.

## 1 Introduction

Chemotherapy is a standard first-line drug treatment of cancer that travels through the body and kills fast-growing cells including cancer, normal, and healthy cells. Damage to healthy cells carries a risk of side effects such as nausea, vomiting, anorexia, muscle weakness, and fatigue. Muscle tissue in particular is susceptible to off-target chemotherapy effects due to its dense nucleation and reliance on cell turnover. Depending on the type of cancer, location, drug(s), dose, and the patient’s general level of health, the chemotherapy side effects can vary person to person. While some side effects are mild and treatable, others can cause serious complications. Evidence suggests that chemotherapeutic agents, including cyclophosphamide, 5-fluorouracil (5-FU), cisplatin, and methotrexate, regardless of their effects on tumour growth can induce unintended skeletal muscle and adipose tissue wasting which is called cachexia [1–3].

Cancer cachexia is a debilitating condition correlated with several types of malignant cancers and is associated with poor response to treatment and decreased survival rates [4–9]. Since cachexia is a common side effect of cancer and to chemotherapy treatment, one may wonder whether or not the activated molecular mechanisms that induce muscle loss are similar.

Despite many recent studies, the underlying mechanism of chemotherapy-induced or of cancer-induced cachexia are still only partially understood. Daumrauer *et al*. [3] demonstrated that chemotherapy can promote cachexia by activating a muscle atrophy process. They examined the effect of cisplatin, a standard chemotherapeutic agent, on a colon-26 (C26) murine cancer cachexia model and reported that although cisplatin is able to strongly reduce tumour burden, it can also promote muscle atrophy through the activation of the neuclear factor *κ* B (NF-*κ*B) pathway. Further, in [10], it was shown that NF-*κ*B targets not only muscle fibers but also muscle stem cells. During cancer cachexia, in the muscle microenvironment, additional activation of NF-*κ*B leads to Pax7 deregulation, which impairs myogenic cell differentiation [10]. In [11], the authors compared differentially expressed muscle proteins from exposure to either the C26 cancer cell line or the chemotherapy agent folfiri (a combination of 5-fluorouracil, leucovorin, and irinotecan). With 240 commonly modulated proteins, they suggested that cancer and chemotherapy contribute to muscle loss by activating common signalling pathways. Their analysis also determined that mitochondrial dysfunctions were present in both experimental conditions. Later, they reported in [12,13] that inhibition of the activin receptor 2B (ACVR2B) signalling represents a potential therapeutic strategy to prevent muscle loss following folfri-treatment. Their results were consistent with previous observations reporting beneficial effects of the inhibitor ACVR2B/Fc in cachexia treatment in a C26 mouse model [14].

Despite these similarities, emerging research suggests that chemotherapy-induced muscle loss differs from cancer-induced loss [15], and that the dysfunction may persist for years after treatment [16]. Chemotherapy can lead to increased mitochondrial reactive oxygen species (mtROS) and dysfunction that leads to fatigue, muscle loss, impaired regenerative capacity, pain and exercise intolerance [17]. Mitochondrial toxicity by cancer [18, 19] or chemotherapeutic agents [20–22] is thus another possible mechanism of cachexia. In fact, it was shown that the metabolic perturbations from cancer-induced or chemotherapy-induced cachexia are quite distinct, with significant differences in amino acid catabolism, lipoprotein metabolism, and inflammation [15]. The review [17] summarizes the molecular origins of chemotherapy-induced myopathy with a focus on the mitochondria. They present a hypothetical model proposing that chemotherapy administration to healthy skeletal muscle affects nuclear DNA and mitochondrial DNA (mtDNA) resulting in dysfunction and increased mtROS leading to satellite cell replicative failure, impaired regenerative ability, and progressive muscle loss.

In addition to being the essential energy producers in cells, recent studies have identified a direct role for mitochondrial activity in the regulation of stem cell fate and differentiation [23–27]. The mitochondria has an important role in the metabolism of lipids, carbohydrates, and proteins and acts as a center for coordinating extrinsic and intrinsic signals to direct growth, proliferation, differentiation, and cell death [28, 29]. Mitochondrial stress and dysfunction plays a key role in aging and apoptosis [30,31], leads to a wide range of diseases [32,33], and has been implicated as a major player in the health hazards of spaceflight [34].

In this paper we mathematically investigate cachexia due to the administration of chemotherapy. Experimental data for daily and 5-days-a-week schedules and several doses of 5-FU are used to parameterize the model. The anti-metabolite 5-FU chemotherapy agent is a commonly used treatment for several cancers including colorectal, pancreatic, head and neck, breast, and cancers of the aerodigestive tract [35]. In previous work, [36], we developed a mathematical model of muscle tissue growth and homeostasis in a healthy state and in a cancer-induced cachectic state, with and without treatment by an ACVR2B inhibitor. Here, we modify our healthy muscle tissue model to investigate the effects of chemotherapy on satellite and muscle cells. We aim to better understand the mechanism of chemotherapy-induced cachexia as well as to explore alternate dose schedules that preserve lean mass and thus reduce treatment side effects and improve patient quality of life.

This paper is organized as follows: in Section 2, we examine the experimental data from 5-FU administration in mice under various dosages and two different dosing schedules. We then construct our mathematical model for chemotherapy-induced cachexia. In Section 3, we discuss model parameterization, fit two model parameters that cannot be extracted from the literature, and discuss the model sensitivity of these two parameters. In Section 4, we use the model to examine alternate dosing schedules and discuss metrics to compare the therapeutic efficacy of treatments considering both lean mass preservation and tumour control. We then explore aging effects and morphine use in chemotherapy-induced cachexia. The paper concludes with a discussion in Section 5.

## 2 Model Development

In this section we introduce the experimental data and then construct a mathematical model for chemotherapy-induced muscle wasting.

### 2.1 Dose-Dependence of Chemotherapy-Induced Muscle Loss

Makino *et al*. [37] reported a comparative study between daily and 5-days-a-week oral administration of 5-FU in 6 week old CDF1 male mice. They found that body weight decreased under drug administration in a nonlinear dose-dependent manner. We extrapolated the lean mass from these body weight measurements by assuming an initial weight of 25 g and then assuming the body composition was approximately 40% lean mass. Our estimates of the lean mass response to F-5U administration following a daily or 5-on, 2-off schedule are shown in Figure 1. The dosing schedule lasted 4 weeks and the mice were observed for a further 4 weeks. The two regimes administered either 28 doses following the daily schedule, or 20 doses following the 5-on, 2-off schedule over experimental days 0 to 27. Various dosages were tested ranging from 14 − 35 mg/kg for the daily schedule and 24 − 60 mg/kg for the 5-on, 2-off schedule. Mortality was reported for the daily schedule in 4/5 mice receiving 35 mg/kg. For the 5-on, 2-off schedule, mortality was reported in 5/5 mice receiving 60 mg/kg and 3/5 mice receiving 50 mg/kg.

**Figure 1:**
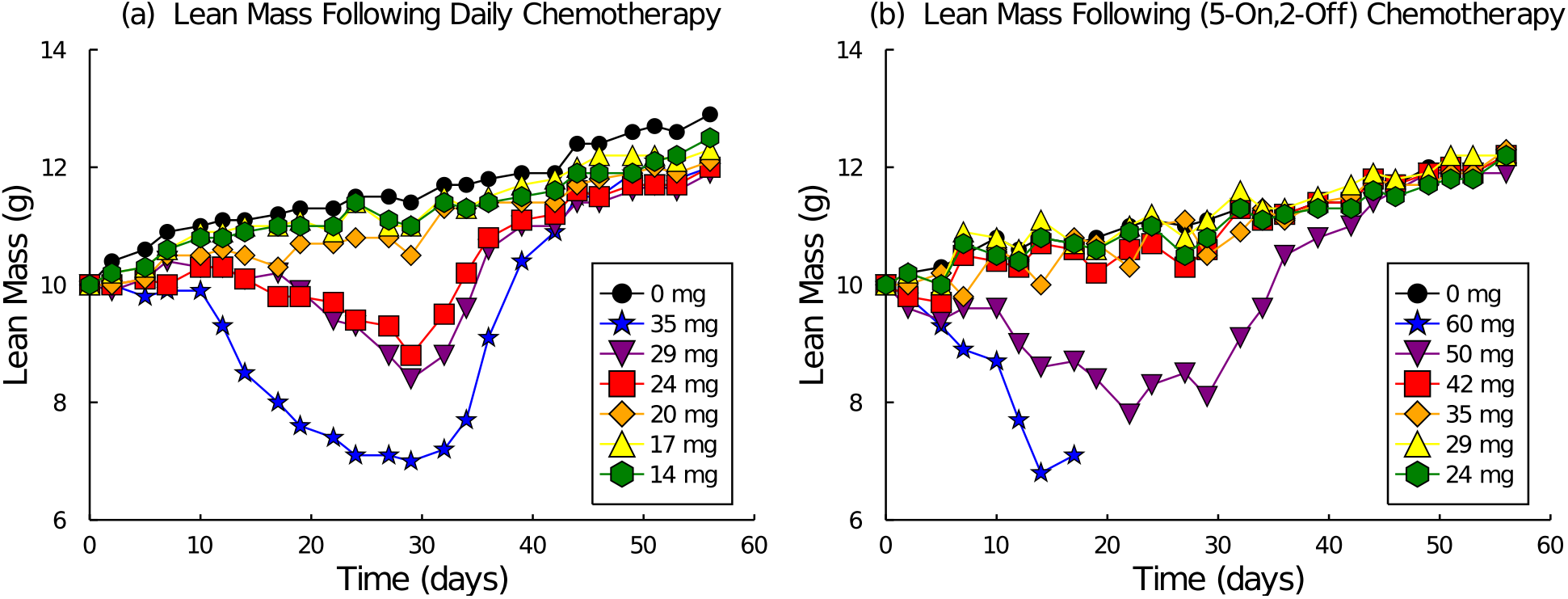
Lean mass response to 5-FU chemotherapy administration estimated from experimental data [37] following either a daily schedule (a) or a 5-on, 2-off schedule (b) over days (0, 29).

The experimental data contains some unique features of the dose response. First consider the daily schedule shown in Figure 1(a). Note that there is a striking lag in mass lost over the first 10 days. This lag in effect is not a true delay of response, however, because there is no lag in recovery when the treatment is stopped on day 28. Note also that doses 14, 17, and 20 mg/kg track approximately with the control data, doses 24 and 29 mg/kg demonstrate some loss, and dose 35 mg/kg demonstrates significant loss and an accelerated rate. The spread of these curves is not linear, meaning the amount of mass lost does not increase linearly with dose. In fact, this data suggests a strong nonlinear dose-dependence.

Upon examination of the 5-on, 2-off schedule, see Figure 1(b), we note the slight oscillations in lean mass due to the drug holiday every 5 days. Larger doses (greater than 42 mg/kg) no longer seem to have a 10-day lag before noticeable loss. Doses less than 42 mg/kg seem to all approximately track with the control, indicating no significant mass loss. Of interest is that after 28 days of treatment, the 50 mg/kg group, which lost considerable mass, recovers quite quickly and immediately with 2/5 mice surviving to the end of the experiment, whereas the 60 mg/kg group did not last the full experiment.

### 2.2 Mathematical Model

We now shift to the development of a mathematical model for chemotherapy-induced cachexia. To begin, we introduce a pharmacokinetic model for chemotherapy and then introduce a model of healthy muscle tissue based on stem cell lineage dynamics. These models are coupled together to describe the muscle loss due to chemotherapy in a manner that captures the unique features observed in the data. Finally we add a tumour model in order to assess the efficacy of treatment plans on both tumour control and muscle preservation.

#### 2.2.1 Pharmacokinetic Model of Chemotherapy

Compartment models are often used to describe the kinetic behaviour of drug concentration profiles in the body. We use a 2-compartment model (model 7 of Win-NonLin [38]) describing a bolus input of the drug, transit between the two compartments, and first order clearance. This models the kinetic behaviour of the chemotherapy drug concentration in the plasma, *C*_1_(*t*), and tissue, *C*_2_(*t*), compartments [39, 40]. The pharmacokinetic equations are:

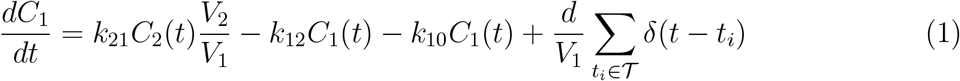

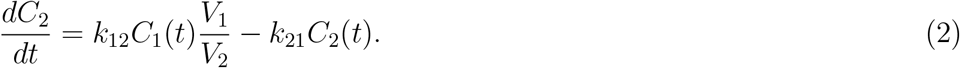

Drug concentrations *C*_1_ and *C*_2_ are measured in *μ*g/ml for each compartment. *V*_1_ and *V*_2_ are the corresponding distributed volumes of the drug after one dose in each compartment (in ml/kg). The drug dosage *d* is measured in *μ*g/kg and administered instantaneously over the set of treatment times *t*_*i*_ ∈ *𝒯* by the dirac-delta train Σ*δ*(*t − t*_*i*_). The rate constants *k*_12_ and *k*_21_ (in days^−1^) describe the transfer of drug between the plasma and tissue compartments. The drug clearance rate is described by *k*_10_ in days^−1^.

To simulate the dosing schedule, we interrupt the computational simulation at each treatment time *t*_*i*_ ∈ *𝒯* and add another dose amount 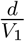 to the solution *C*_1_(*t*_*i*_). This is done via a *discrete call back* to our numeric solver in the Julia programming language [41]. The daily treatment schedule administers 28 doses, one on each day in 𝒯_*daily*_ = [0, 1, 2, …, 27]. The (5-on, 2-off) treatment schedule administers 20 doses over the first 28 days, following the schedule *𝒯*_(5,2)_ = [0, …, 4, 7, …, 11, 14, …, 18, 21, …, 25].

#### 2.2.2 Model of Healthy Muscle Tissue

In previous work [36] we introduced a stem-cell lineage model for muscle tissue. Let *S* describe the satellite (stem) cell compartment and *M* the differentiated muscle fibre com-partments as volumes measured in mm^3^. The stem cells are assumed to divide symmetrically to produce either two stem cells with probability *p*, or two differentiated muscle cells with probability 1 *− p*. The model equations for a healthy muscle tissue are thus:

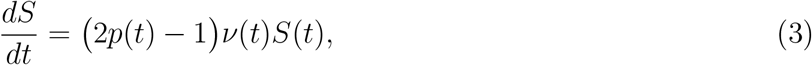

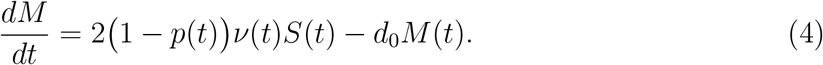

Here *p*(*t*) is the probability of satellite cell self-renewing division, *ν*(*t*) is the proliferation rate, and *d*_0_ is the natural muscle cell death rate.

Feedback from the muscle compartment to the satellite cell compartment is required to regulate tissue growth and regeneration by decreasing the probability of self-renewal for large muscle sizes. We describe this feedback mathematically as:

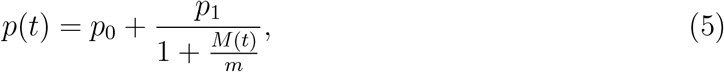

where *p*_0_ is the base probability and *p*_1_ is a perturbation probability activated in early growth or after muscle damage. Parameter *m* is a half-saturation coefficient for the feedback dynamic of muscle mass *M*(*t*).

The satellite cell proliferation rate, *ν*(*t*), also has a feedback mechanism that reduces proliferation as the muscle nears its equilibrium size:

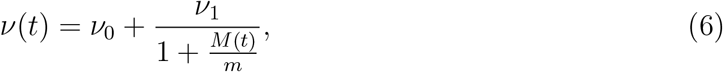

where *ν*_0_ is the base proliferation rate and *ν*_1_ is a perturbation proliferation rate activated in early growth and after tissue damage. Parameter *m* is again a half-saturation coefficient for the feedback dynamic of muscle mass *M*(*t*), assumed to be the same as in the probability *p*(*t*) for simplicity.

In order to compare our model to lean mass measurements in grams, we convert these volumetric predictions to mass via the formula: lean mass = 0.002(*S* +*M*) g. The conversion factor of 0.002 g/mm^3^ was estimated in [36].

#### 2.2.3 Model of Chemotherapy-Induced Cachexia

The biological mechanisms of muscle loss due to chemotherapy are not well understood. We explored several mathematical forms that prescribed increased death rates in both model compartments due to chemotherapy, but none of these models provided a good fit to the experimental data. With increased death added to the model, the recovery of lost mass upon cessation of treatment was accompanied by very large overshoot (well above the control). This overshoot is due to a transient disruption to the stem cell ratio, and hence accelerated regrowth. The feedback mechanism from lost muscle cells stimulates proliferation of stem cells, allowing for the muscle recovery, but also for the overshoot. It is important to note that in our previous work, [36], cancer-induced cachexia was partially described mathematically by increased stem, *S*(*t*), and muscle cell, *M*(*t*), death rates. The fact that the same model formulation does not fit the experimental data here, suggests that chemotherapy-induced cachexia is activating a different biological mechanism than we observed in cancer-induced cachexia.

Off-target chemotherapy effects may cause skeletal muscle damage and loss by inducing mitochondrial toxicity and dysfunction [17, 18, 42–46]. Mitochondrial metabolism is now known to be a key regulator of stem cell activation, fate, and differentiation [25, 26]. Furthermore, mitochondrial dysfunction can lead to senescence and stem cell aging [26]. Thus, we can explore potential dysfunction of our stem-cell lineage model of muscle tissue due to the applied chemotherapy’s effects on mitochondria and thus stem cell regulation.

As a simple first assumption, we assume that chemotherapy can slow the successful proliferation rate of stem cells. Biologically this mimics the effects of dysregulated mitochondria (and thus the dysregulated stem cell actions) of cells exposed to 5-FU. Further, if the differentiation program is disrupted by the drug, then progenitor cells will die out rather than form new muscle cells, impairing the regeneration cascade. We model these mechanisms of mass loss as a reduction in successful proliferation and differentiation, and propose the following modification to the muscle tissue model:

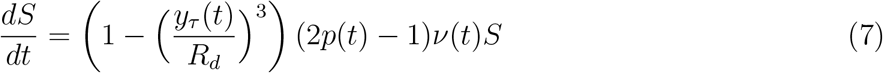

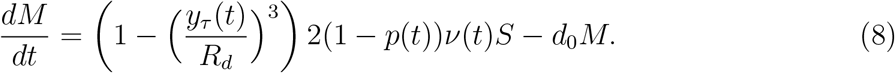

The chemotherapy effect is included via a cubic function of a moving time-averaged exposure function *y*_*τ*_ (*t*). Parameter *R*_*d*_ (in *μ*g/ml) is a thresholding constant to scale the cubic function. The average expose *y*_*τ*_ is defined as

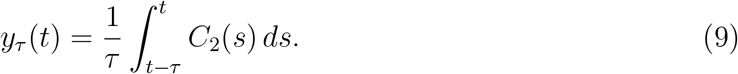

This *τ* -average exposure of the tissue to drug concentration *C*_2_(*t*) is assumed to cause the disruption to muscle homeostasis. The average over *τ* -days is used to slow the drug’s response dynamics and to capture the initial delay in mass loss observed under the daily schedule. The cubic functions in equations (7) and (8) reflect the non-linear dose response of the lean mass to treatment. Maintaining a balance between stem and muscle cells is key to controlling overshoot of the control upon recovery. The symmetric nature of these equations also minimizes disruption of the stem cell ratio under chemotherapy.

#### 2.2.4 Chemotherapy Effect on Tumour Burden

Lastly, we explore the effect of various dosing schedules on a simulated tumour burden. For this we assume an exponential-linear tumour growth model [39] and use parameter values previously fit to C26 murine carcinoma experimental data [36]. The tumour growth equation is

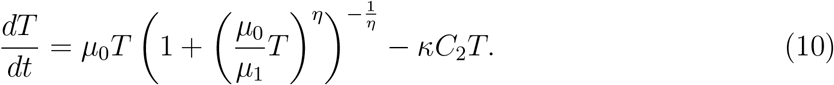

Here *η* = 20 controls the transition from the exponential growth phase to the linear growth phase, and is sufficiently large enough to make the transition near-instantaneous. Parameter *μ* = 0.446 days^−1^ is the exponential growth rate, and *μ*_1_ = 116.0 mm^3^ days^−1^ is the linear growth rate [36]. We assume an initial condition of *T*(0) = 10 mm^3^. The chemotherapy effect in the second term on the right-hand-side assumes cell death is proportional to the drug concentration within the tissue, *C*_2_(*t*) and the tumour volume *T*(*t*). We choose a drug efficacy of *κ* = 0.13 ml *μ*g^−1^ day^−1^.

## 3 Model Parameterization

The healthy muscle tissue model was previously parameterized in [36] by fitting model equations (3) and (4) to an extended growth curve of US-bred CDF1 male mice [47]. The pharmacokinetic model parameters for 5-FU were reported in [39, 40]. The exponential-linear tumour growth parameters were reported in [36] after fitting to experimental data. This leaves *R*_*d*_, *τ*, and *κ* as the only unknown model parameters for our model. The chemotherapy efficacy, *κ* was chosen to be 0.13 ml *μ*g^−1^ day^−1^ to ensure certain treatment schedules achieved a clearly reduced tumour burden. This leaves only *R*_*d*_ and *τ* as parameters to fit for this chemotherapy-induced cachexia model. All parameter values reported in the above references are listed in Table 1.

**Table 1:** Model parameters and their estimated values from previous data fitting reported in the cited reference.

| Description | Value | Reference |
| --- | --- | --- |
| drug clearance rate from plasma | $k_{10} = 151.2 \text{ days}^{-1}$ | [39] |
| transfer rate from plasma to tissue | $k_{12} = 5.62 \text{ days}^{-1}$ | [39] |
| transfer rate from tissue to plasma | $k_{21} = 2.31 \text{ days}^{-1}$ | [39] |
| distribution volume of plasma | $V_1 = 0.71 \times 10^3 \text{ ml/kg}$ | [39] |
| distribution volume of tissue | $V_2 = 0.1 \times 10^3 \text{ ml/kg}$ | [39, 40] |
| homeostatic probability of self-renewal | $p_0 = 0.479$ | [36] |
| maximum perturbation to probability | $p_1 = 0.133$ | [36] |
| homeostatic proliferation rate | $\nu_0 = 0.087 \text{ days}^{-1}$ | [36] |
| maximum perturbation to proliferation | $\nu_1 = 5.591 \text{ days}^{-1}$ | [36] |
| natural muscle fiber death rate | $d_0 = 0.05 \text{ days}^{-1}$ | [36] |
| half-saturation coefficient for feedback | $m = 1000 \text{ mm}^3$ | [36] |
| exponential tumour growth rate | $\mu_0 = 0.446 \text{ days}^{-1}$ | [36] |
| linear tumour growth rate | $\mu_1 = 116.0 \text{ mm}^3 \text{ days}^{-1}$ | [36] |
| transition factor from exponential to linear growth | $\eta = 20$ | [39] |

### 3.1 Drug Pharmacokinetics and Average Exposure Function

To demonstrate the effect of the *τ* -day averaging function *y*_*τ*_ (*t*) on chemotherapy exposure, we simulate the two dosing schedules for a dose of 35 mg/kg, Figure 2. The drug kinetics following one dose are shown in Figure 2[a]. The daily schedule and resulting 5-day average exposure are shown in Figure 2[b] and [c], respectively. and the (5-on, 2-off) schedule and resulting 5-day average exposure are shown in Figure 2[d] and [e], respectively.

**Figure 2:**
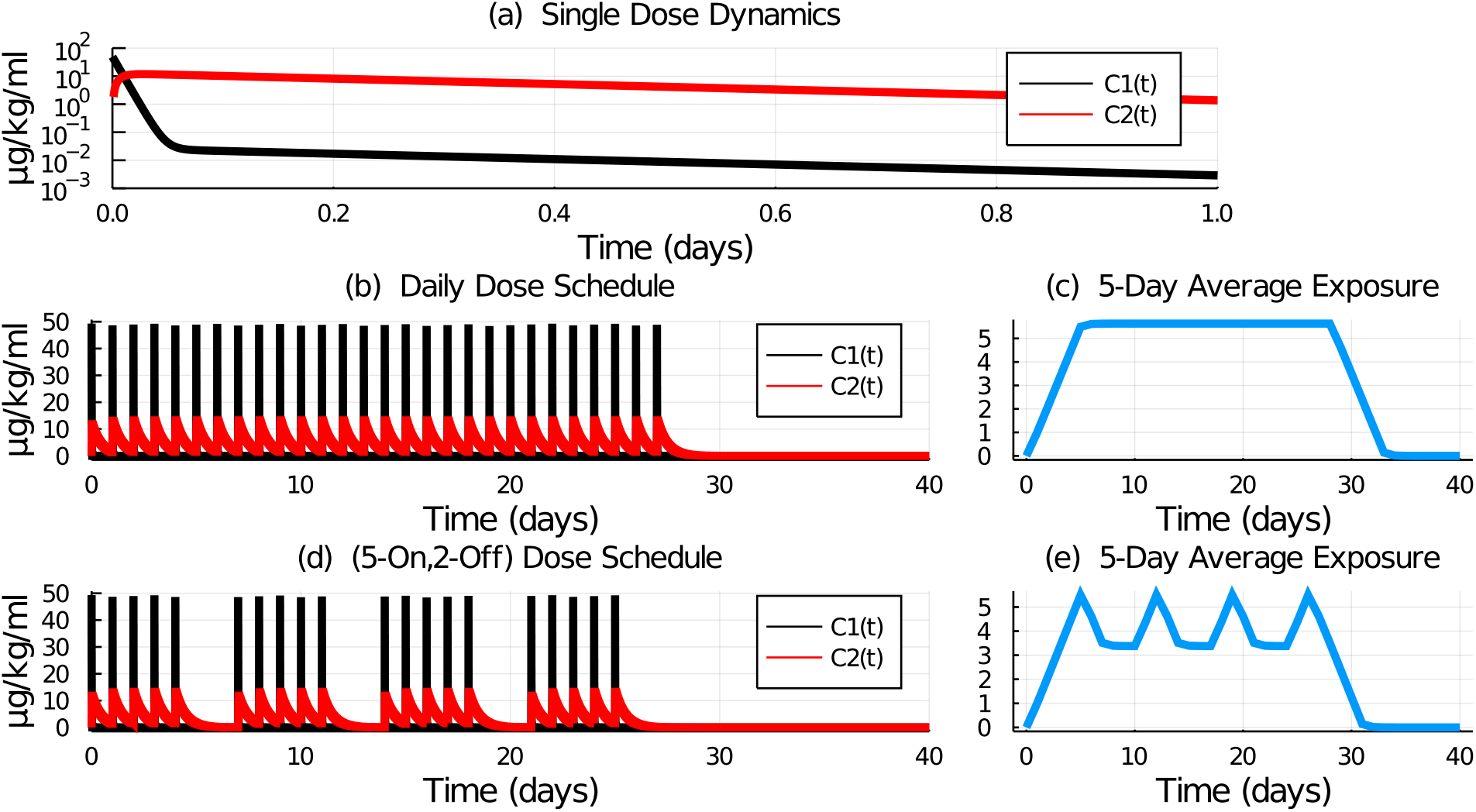
Model simulation of plasma and tissue concentrations of 5-FU given (a) one dose, (b) a sequence of 28 daily doses, and (d) a sequence of 20 doses following the (5-on, 2-off) schedule. The dose of 35 mg/kg was simulated by numerically solving equations (1) and (2). The 5-day average exposure for (c) the daily schedule and (e) the (5-on, 2-off) schedule was computed from equation (9) for *τ* = 5.

### 3.2 Parameter Fitting to the Experimental Data

It remains to determine parameters *R*_*d*_ and *τ*. We run our simulations assuming an initial lean mass of 10 g, corresponding to 6-week old mice: this gives initial conditions of *S*(0) = 267.5 mm^3^ and *M*(0) = 4732.5 mm^3^ (see [36][equation 8] for method to estimate initial volumes based on the approximate stem cell ratio of certain aged mice).

We aim to find global *R*_*d*_ and *τ* values that enable our model to best fit the experimental data for both dosing schedules and all dose curves. To find the value of *R*_*d*_ and *τ* we fit the model prediction for lean mass to the experimental data using the sum of squared errors as the optimization function. To be clear, the cost function used is the total sum of all squared deviations between each non-zero dose curve and its model prediction, so as to fit all dose curves at once. A hybrid approach is used for this data fitting problem. First, *τ* is fixed and *R*_*d*_ is fit using a grid search method. Then the value of *τ* is increased and the process is repeated. Integer values of *τ* were tested over [1, 15] days.

The best fit for each dose schedule is shown in Figure 3. The daily schedule has a minimum SSE when *τ* = 11 and *R*_*d*_ = 6.18. The long value of *τ* here helps the model capture the lag in mass loss upon initiation of daily treatment. The (5-on, 2-off) schedule has a minimum SSE when *τ* = 5 and *R*_*d*_ = 7.86. The short value of *τ* here helps the model capture the oscillations in mass loss due to the drug holiday over weekends. Figure 3 also shows the average exposure function and the stem cell ratio (*S/M*) of the tissue. The treatment disrupts the normal tissue stem ratio of approximately 5% as the stem compartment is activated by the feedback mechanisms in order to replenish the lost mass. This disruption is transient.

**Figure 3:**
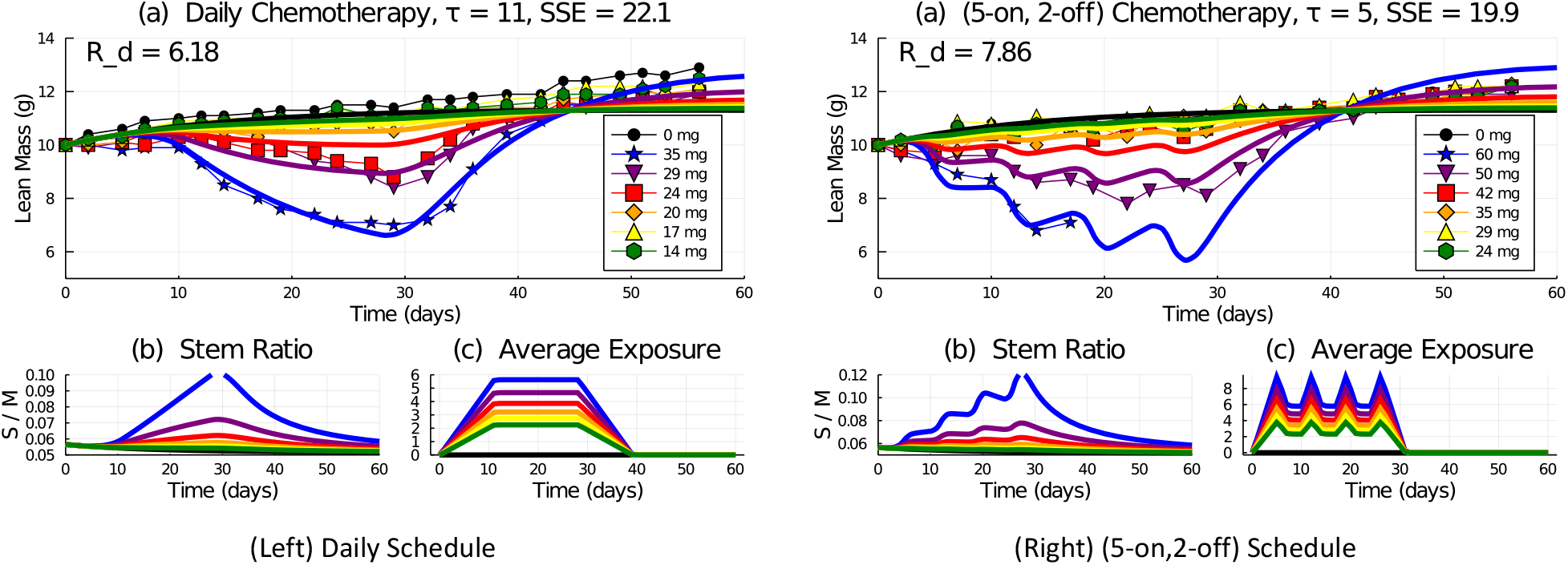
Model simulation with fitted parameters *R*_*d*_ and *τ* for the (left) daily and (right) (5-on, 2-off) chemotherapy schedules administering various doses over days (0, 27). Top: model simulation of lean mass response to chemotherapy. Bottom: model predicted stem cell ratio (b) and average exposure function *y*_*τ*_ (*t*) (c).

Finally, we plot the SSE for all values of *τ* and both dosing schedules in order to determine global values we can apply for any dosing schedule. By examination of Figure 4(a), we select *τ* = 8 as our global value as it is the mid-point between the minimum SSEs for each schedule (daily schedule has minimum SSE at *τ* = 5 and (5-on, 2-off) schedule has minimum SSE at *τ* = 11). The global value of *R*_*d*_ is then chosen as the average of the fitted *R*_*d*_ values for each schedule when *τ* = 8, giving *R*_*d*_ = 6.8, see Figure 4(b). We thus use values of *τ* = 8 and *R*_*d*_ = 6.8 as our global parameters for any dosing schedule and/or applied dose.

**Figure 4:**
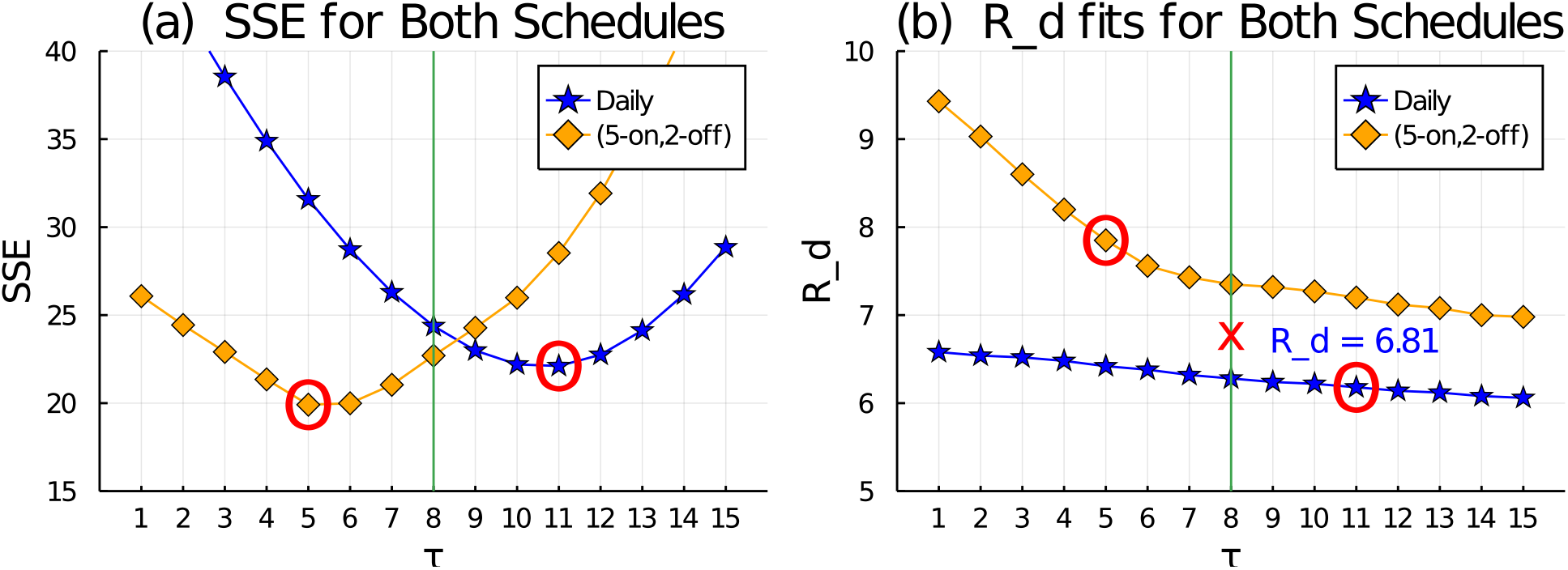
The sum of squared errors (SSE) (a) and best-fit *R*_*d*_ value (b) for all tested values of *τ* over [1, 15] and both dosing schedules. From (a), the SSE is minimum when *τ* = 11 for the daily schedule and *τ* = 5 in the (5-on, 2-off) schedule. Hence we take the optimal value as the mid-point at *τ* = 8. From (b), when *τ* = 8, the approximate average of the two fitted *R*_*d*_ values is *R*_*d*_ = 6.8.

### 3.3 Model Sensitivity Analysis

To explore the sensitivity of the two new model parameters *R*_*d*_ and *τ*, we computed dynamic sensitivity analyses. First, we found the resulting lean mass response following a 10% increase or decrease to parameters *R*_*d*_ = 6.8 or *τ* = 8. Then we computed the sensitivity coefficients and relative sensitivity coefficients. For parameter *ρ* ∈ {*R*_*d*_, *τ*} and at time *t*_*i*_, the sensitivity coefficient of the lean mass (*LM*) computation is:

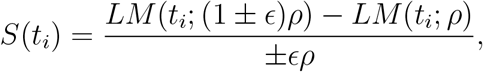

and the relative sensitivity coefficient is

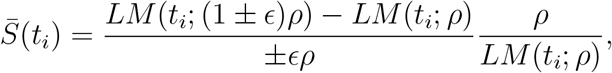

where *ϵ* = 10^−5^ is a small perturbation parameter, small enough that the coefficients stabilize in their values. These computations were performed during either a daily or (5-on, 2-off) chemotherapy schedule over the first 28 days, as in the experimental data. The analysis was carried out further to explore the model response as it returns to steady-state following the treatment perturbations.

The parameter sensitivity for *R*_*d*_ is shown in Figure 5. A 10% change to *R*_*d*_ causes a larger perturbation to the solution for the daily schedule than the 5-day schedule, because the daily schedule applies 28 doses whereas the 5-day schedule only applies 20 doses. The 2-day holidays also reduce the cumulative effects of the perturbation. The dosage difference between the two schedules is also apparent in the sensitivity coefficients, with the daily schedule accumulating a larger sensitivity to parameter *R*_*d*_ with each successive dose. In all instances, the effects of the parameter perturbations are transient and decay once treatment is stopped.

**Figure 5:**
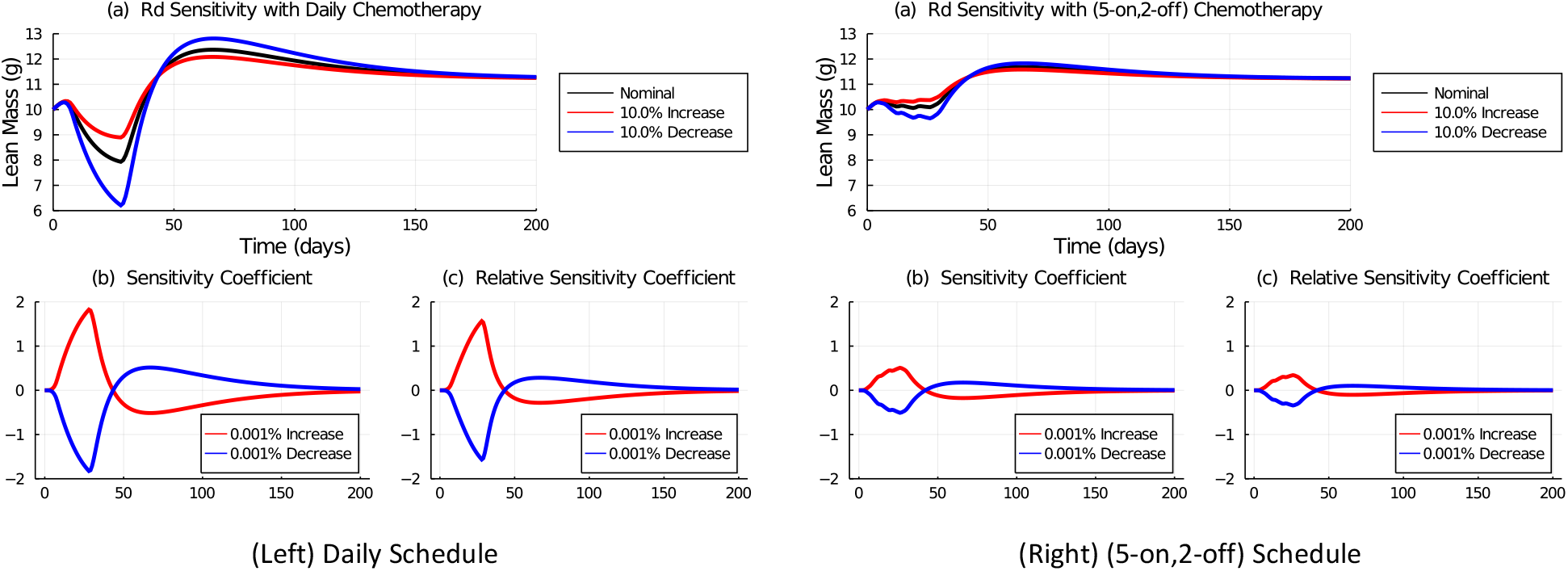
Parameter sensitivity analysis for parameter *R*_*d*_ using a daily (Left) or (5-on, 2-off) (Right) chemotherapy schedule. The nominal value is *R*_*d*_ = 6.8. Top row: (a) the value of *R*_*d*_ is increased or decreased by 10% and the resulting lean mass is shown. Bottom row: the sensitivity coefficient (b) and relative sensitivity coefficient (c) are computed with a 0.001% change to the nominal value. Chemotherapy is applied over the first 28 days.

Figure 6 shows the sensitivity analysis of parameter *τ*. The model is less sensitive to changes in *τ* than *R*_*d*_, as *τ* is related to the time-latency of the chemotherapy effect, rather than the strength of the drug-induced mass loss for *R*_*d*_. A 10% change in *τ* does not significantly perturb the solution, and the sensitivity coefficients are small in magnitude and quickly decay once treatment is stopped.

**Figure 6:**
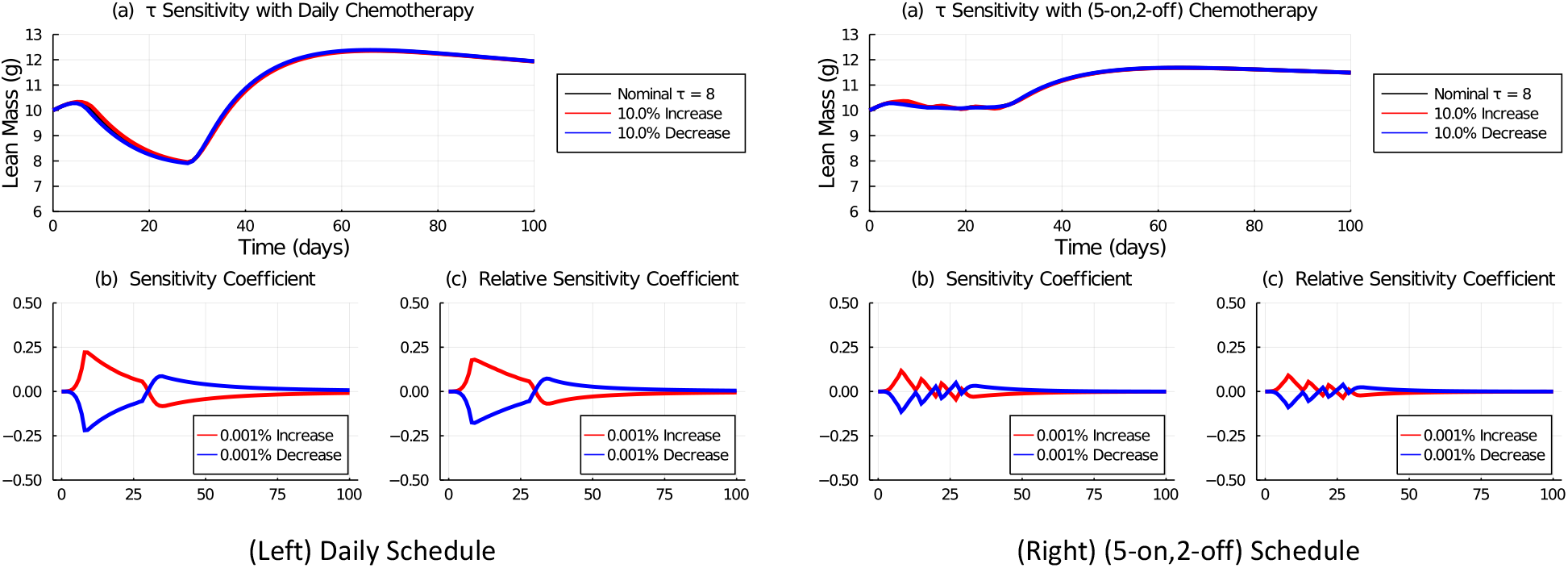
Parameter sensitivity analysis for parameter *τ* using a daily (left) or (5-on, 2-off) (right) chemotherapy schedule. The nominal value is *τ* = 8. Top row: the value of *τ* is increased or decreased by 10% and the resulting lean mass is shown. Bottom row: the sensitivity coefficient (b) and relative sensitivity coefficient (c) are computed with a 0.001% change to the nominal value. Chemotherapy is applied over the first 28 days.

## 4 Results

Given that chemotherapy with 5-FU can cause muscle loss, we now use our mathematical model to explore potential cancer treatment strategies that aim to minimize this loss while simultaneously treating a tumour, and thus improve patient quality of life.

### A Standard Weekly Dose

One way to classify chemotherapy schedules is to compare the total weekly dose. Following [37], we define our standard daily schedule as 28 doses of 24 mg/kg per dose giving 168 mg/kg per week, and we define our standard (5-on, 2-off) schedule as 20 doses of 35 mg/kg per dose giving 175 mg/kg per week. We thus aim to minimize muscle loss while maintaining the total weekly dose between a standard daily and standard 5-day schedule (168–175 mg/kg/week).

First, we consider weekly-periodic regimes delivering equal-sized doses. To do this, we test the schedules listed in Table 2. To determine the optimal schedule in terms of muscle mass preservation, we simulate equations (1), (2), (7), and (8) for each dosing schedule. As seen in Figure 7(a), the tested schedules are all approximately equal in maintaining lean mass, with about a 10% drop that is recovered upon treatment termination with overshoot.

**Table 2:** Weekly dosing schedules delivering between 168 - 175 mg/kg/week with equal sized doses.

| Schedule | Dose (mg/kg) | Weekly Total (mg/kg) |
| --- | --- | --- |
| Daily | 24 | 168 |
| (6-on, 1-off) | 28 | 168 |
| (6-on, 1-off) | 29 | 174 |
| (5-on, 2-off) | 34 | 170 |
| (5-on, 2-off) | 35 | 175 |
| (4-on, 3-off) | 42 | 168 |
| (4-on, 3-off) | 43 | 172 |
| (3-on, 4-off) | 56 | 168 |
| (3-on, 4-off) | 57 | 171 |
| (3-on, 4-off) | 58 | 174 |

**Figure 7:**
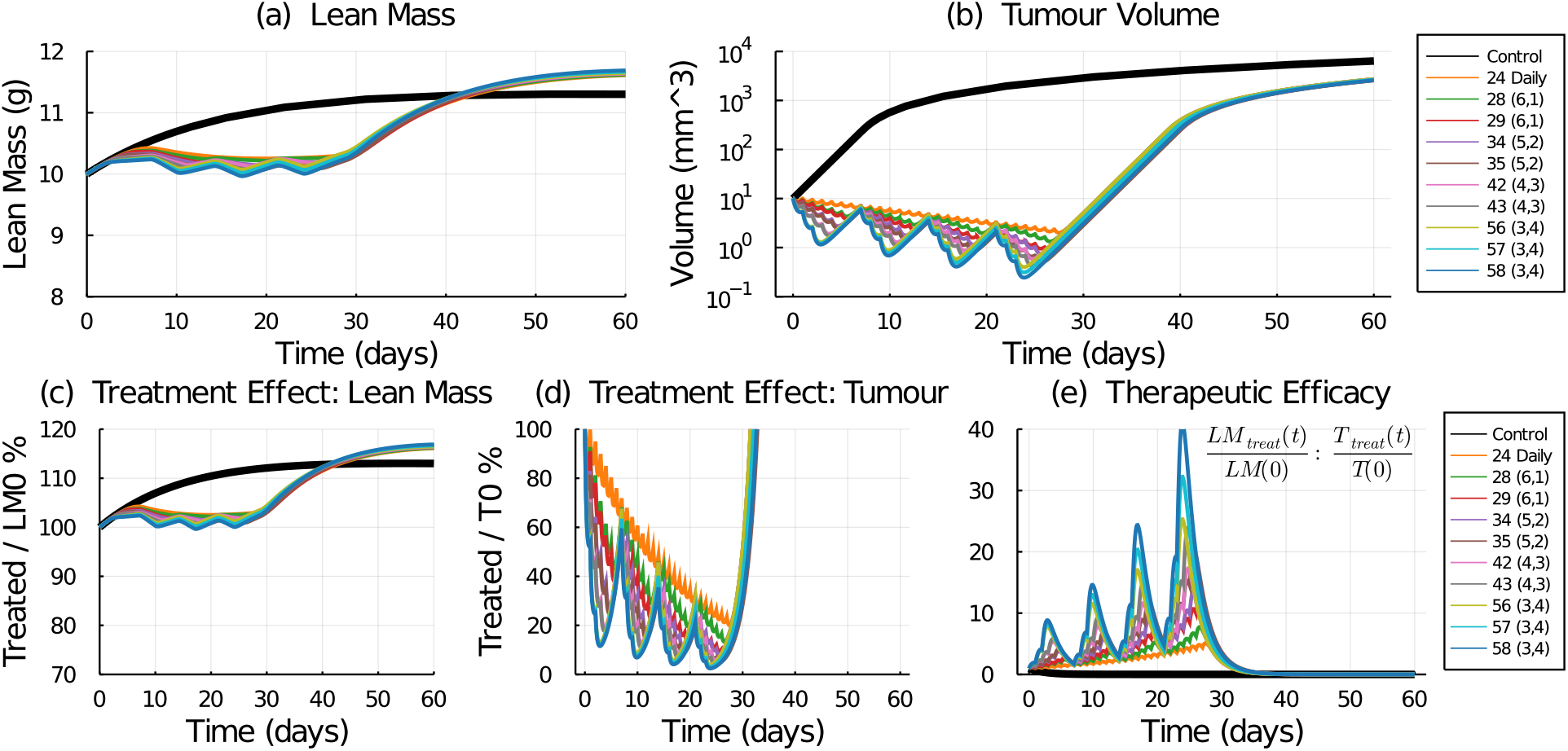
Lean mass (a) and tumour (b) response to treatment schedules listed in Table 2. Treatment effect is computed as the fraction of treated lean mass / control (c), and treated tumour volume / initial tumour volume *T*_0_ (d). Instantaneous therapeutic efficacy is computed by the ratio of the treatment effect fractions (e).

However, simply preserving lean mass is never the objective in cancer treatment - we must consider the treatment effect on a tumour as well. To do this, we simulate the full model (equations (1), (2), (7), (8), and (10)). Note that no cancer-induced cachexia is considered by this model. The effects of the weekly standard treatments (Table 2) applied to a simulated tumour volume with initial size *T*_0_ = 10 mm^3^ are shown in Figure 7(b). Several schedules, including the 5-day schedule successfully reduce the tumour volume below 1 mm^3^, whereas others, including the 24 mg/kg daily schedule, do not. This result is due to the assumption in equation (10) that chemotherapy induces cell death proportional to the drug concentration *C*_2_(*t*).

The largest tumour response is seen with the highest doses: 56–58 mg/kg applied in schedule (3-on, 4-off). In Makino *et al*. [37] they found that the maximum tolerated dose of 5-FU on a daily schedule was 29 mg/kg, and on a 5-day schedule was 42 mg/kg. It is therefore plausible that although it is a higher dose, the 56 − 58 mg/kg doses might be tolerated on a (3-on, 4-off) schedule due to the 4 drug holidays per week. Certainly in the experimental data [37], 50 mg/kg was too high when given 5 days a week as only 1 in 5 mice survived the experiment.

### Therapeutic Efficacy

To assess the treatment’s effect on lean mass, we normalize the model prediction of lean mass response by the initial mass, see Figure 7(c). To assess the tumour response to treatment, we normalize the predicted tumour volume under treatment with the initial tumour size (*T*_0_ = 10 mm^3^ - the volume of the tumour at treatment start), see Figure 7(d). We do not use control (untreated) lean mass or tumour volume since in a clinical setting, only the values upon clinical presentation would be known.

To assess the therapeutic efficacy of each treatment, we examine the ratio of the treatment’s effect on lean mass compared to the treatment’s effect on tumour volume. The formula we use is

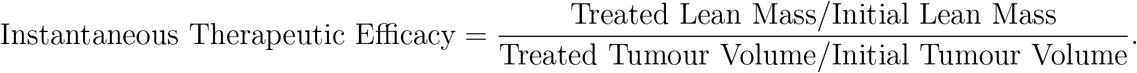

A treatment that preserves lean mass and shrinks the tumour should have a large therapeutic efficacy. The computed therapeutic efficacy for all tested treatments is shown in Figure 7(e). The treatments cause an approximate 10% reduction in lean mass and, depending on the treatment, an approximate 80–95% reduction in tumour volume. In these circumstances, the instantaneous therapeutic efficacy is mostly dominated by the tumour response. This is a consequence of the restriction placed on the tested schedules to deliver a weekly dose within the standard range, and thus affect a consistent but mild decrease in lean mass.

To construct a single numerical value representing the therapeutic efficacy of the treatment, we use the area under the curve (AUC). First, we compute the treatment effects on both the lean mass and tumour volume. The daily schedule administering 24 mg/kg/dose preserves the most lean mass, so it is used to normalize the AUC results. The lean mass AUC ratio for a tested schedule to the 24 mg/kg daily schedule, see Figure 8(a), shows that all tested schedules produce smaller lean mass AUC compared to the 24 mg/kg daily regime. Similarly, we compare the treatment efficacy on the tumour volume to the response from the 24 mg/kg daily standard treatment, see Figure 8(b). Every other tested schedule does a better job at reducing the tumour volume than the daily schedule. Using these two treatment effect ratios, we can then compute the total therapeutic efficacy by comparing them:

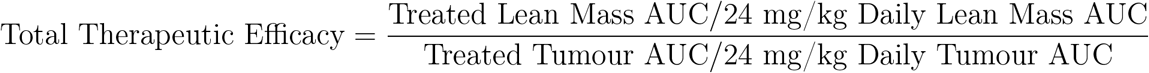

**Figure 8:**
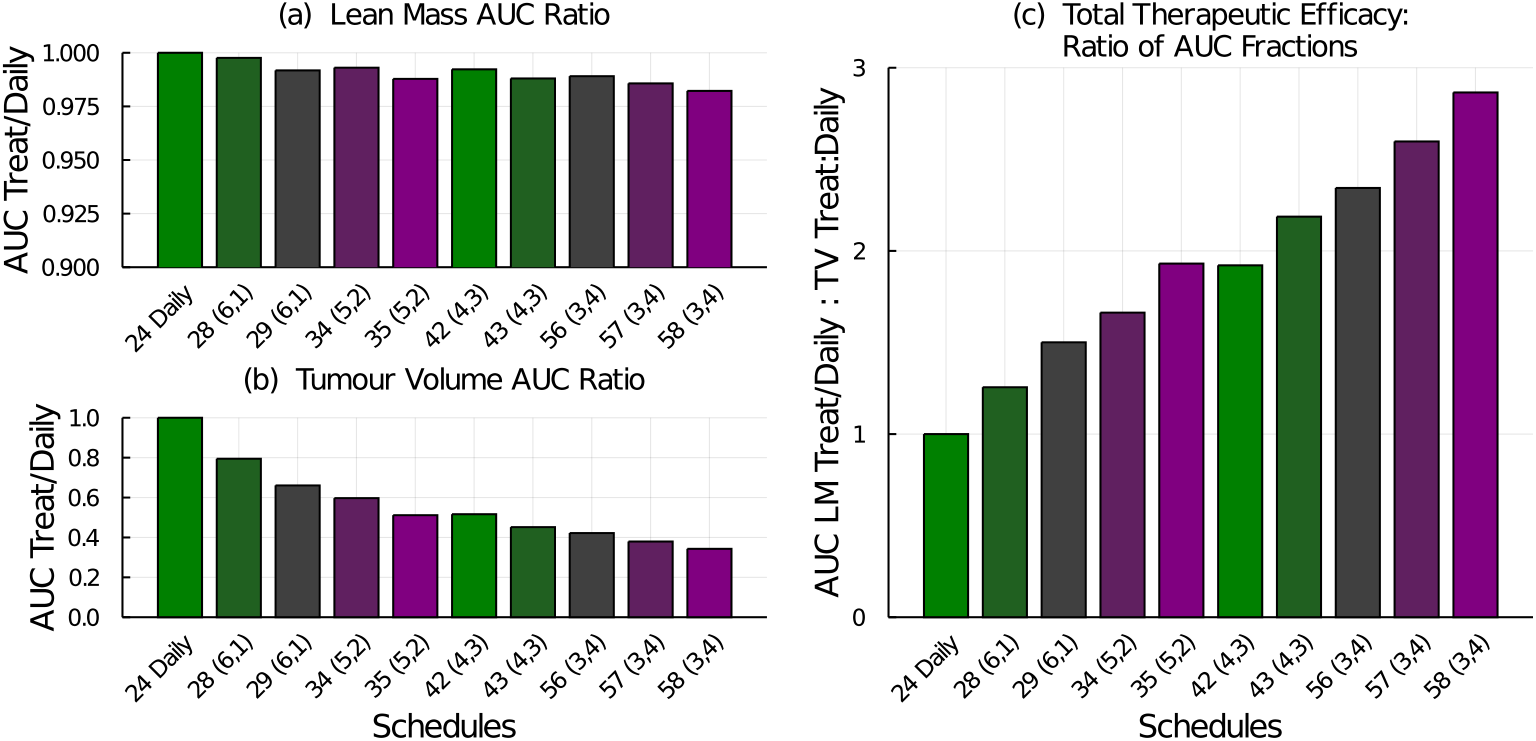
Treatment efficacy for the weekly dose-equivalent chemotherapy schedules listed in Table 2. (a) The AUC ratio for lean mass compares the tested schedule to the 24 mg/kg daily schedule. (b) The AUC ratio for tumour volume compares the tested schedule to the 24 mg/kg daily schedule. (c) The total therapeutic efficacy is the ratio of the lean mass AUC ratio to the tumour volume AUC ratio.

Again, for the tested schedules, since the treatment effect on the tumour volume is more significant than the mass lost, the total therapeutic efficacy metric is dominated by the tumour response. The best treatment in terms of lean mass preservation is the 24 mg/kg daily regime whereas the best treatment in terms of tumour reduction is the 58 mg/kg (3-on, 4-off) regime. The total therapeutic efficacy also suggests that the higher-dosing, less-frequent schedules are preferred over the lower-dosing more-frequent schedules. Interestingly, total therapeutic efficacy is approximately equal for the 35 mg/kg (5-on, 2-off) schedule and the 42 mg/kg (4-on, 3-off) schedule, with the latter having a slight advantage at preserving lean mass.

### Therapeutic Efficacy of Experimental Doses

To examine the robustness of total therapeutic efficacy as a metric to compare treatment schedules, we examine the experimental schedules used in [37] and used for the model parameterization above. Using our mathematical model, we compute the lean mass AUC ratio, tumour volume AUC ratio, and total therapeutic efficacy for the daily and (5-on, 2-off) schedules with various dosages, see Figure 9. From lean mass AUC, it is clear that 60 mg/kg and 50 mg/kg on the (5-on, 2-off) schedule, and 35 mg/kg on the daily schedule result in the most muscle loss, Figure 9(a). These regimes were also toxic and resulted in early mortalities. They also resulted in the most tumour volume reduction predicted by the model, Figure 9(b), and have the highest predicted total therapeutic efficacy, Figure 9(c).

**Figure 9:**
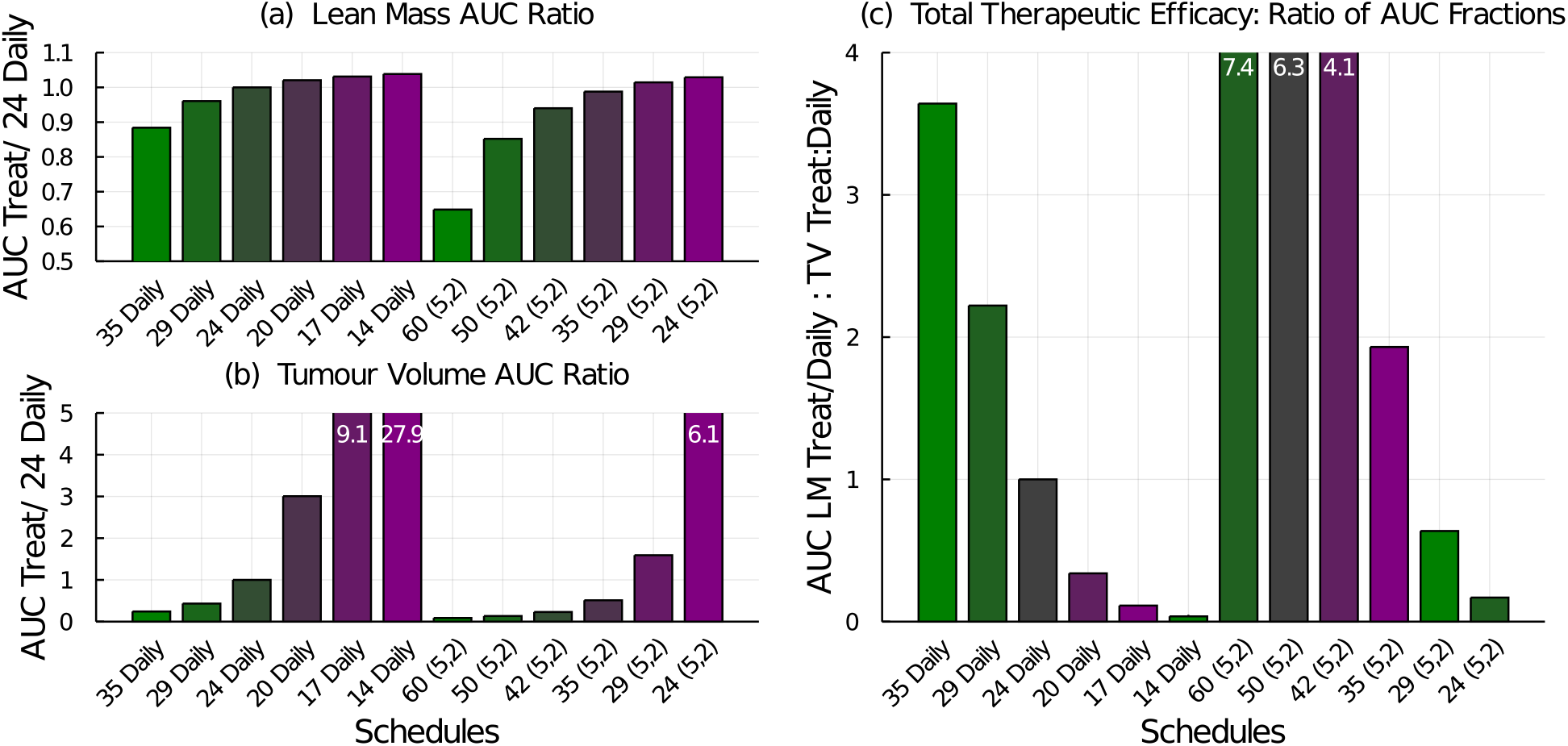
Treatment efficacy for the chemotherapy dosing schedules tested experimentally in [37]. (a) The AUC ratio for lean mass compares the lean mass response from the tested schedule to that of the 24 mg/kg daily schedule. (b) The AUC ratio for tumour volume compares the tumour volume response from the tested schedule to that of the 24 mg/kg daily schedule. (c) The total therapeutic efficacy is the ratio of the lean mass AUC ratio to the tumour volume AUC ratio. If a bar’s height goes above the plot, the height value is written at the top of the bar.

Since the total therapeutic efficacy metric does not take into account drug toxicity, the results must be interpreted carefully. For example, using the lean mass AUC ratio, Figure 9(a), one can eliminate any schedule predicted to cause significantly more lean mass loss than the standard therapy (here the 24 mg/kg daily schedule). This excludes the 35 and 29 mg/kg daily schedules and the 60, 50, and 42 mg/kg (5-on, 2-off) schedules. From the remaining schedules, the total therapeutic efficacy (Figure 9(c)) then shows that the 35 mg/kg (5-on, 2-off) schedule has a bigger therapeutic effect than the standard daily schedule while all other remaining schedules do not perform as well.

### Revisiting a Standard Weekly Dose

Using the AUC ratios and total therapeutic efficacy one can further explore schedules delivering a weekly standard dose. Consider the weekly schedules listed in Table 3 which deliver between 168–175 mg/kg/week. These schedules further break down the dose delivery into weekly patterns that alternate dosing days with drug holidays.

**Table 3:** Dosing schedules that deliver a weekly dose of between 168-175 mg/kg/week. Notation: a schedule described as (4, 1, 1, 1) is a shortening of (4-on, 1-off, 1-on, 1-off), and means doses are given on days 1, 2, 3, 4, and 6 of every week.

| Schedule | Dose (mg/kg) | Weekly Total (mg/kg) |
| --- | --- | --- |
| Daily | 24 | 168 |
| 5 times a week: (5, 2), (4, 1, 1, 1),<br>(3, 1, 2, 1), (2, 1, 3, 1), (1, 1, 4, 1) | 34 | 170 |
| 5 times a week: (5, 2), (4, 1, 1, 1),<br>(3, 1, 2, 1), (2, 1, 3, 1), (1, 1, 4, 1) | 35 | 175 |
| 4 times a week: (4, 3),<br>(3, 2, 1, 1), (2, 2, 2, 1), (1, 2, 3, 1),<br>(1, 1, 1, 1, 1, 1, 1) | 42 | 168 |
| 4 times a week: (4, 3),<br>(3, 2, 1, 1), (2, 2, 2, 1), (1, 2, 3, 1),<br>(1, 1, 1, 1, 1, 1, 1) | 43 | 172 |
| Every Other Day (1, 1) | 48 mg/kg | 168 on average |
| Every Other Day (1, 1) | 50 mg/kg | 175 on average |

The model predicted response of the lean mass, tumour volume, and total therapeutic efficacy are shown in Figure 10. Spreading the doses out across the week preserves lean mass, Figure 10(a), but also decreases the tumour’s response to treatment, Figure 10(b).

**Figure 10:**
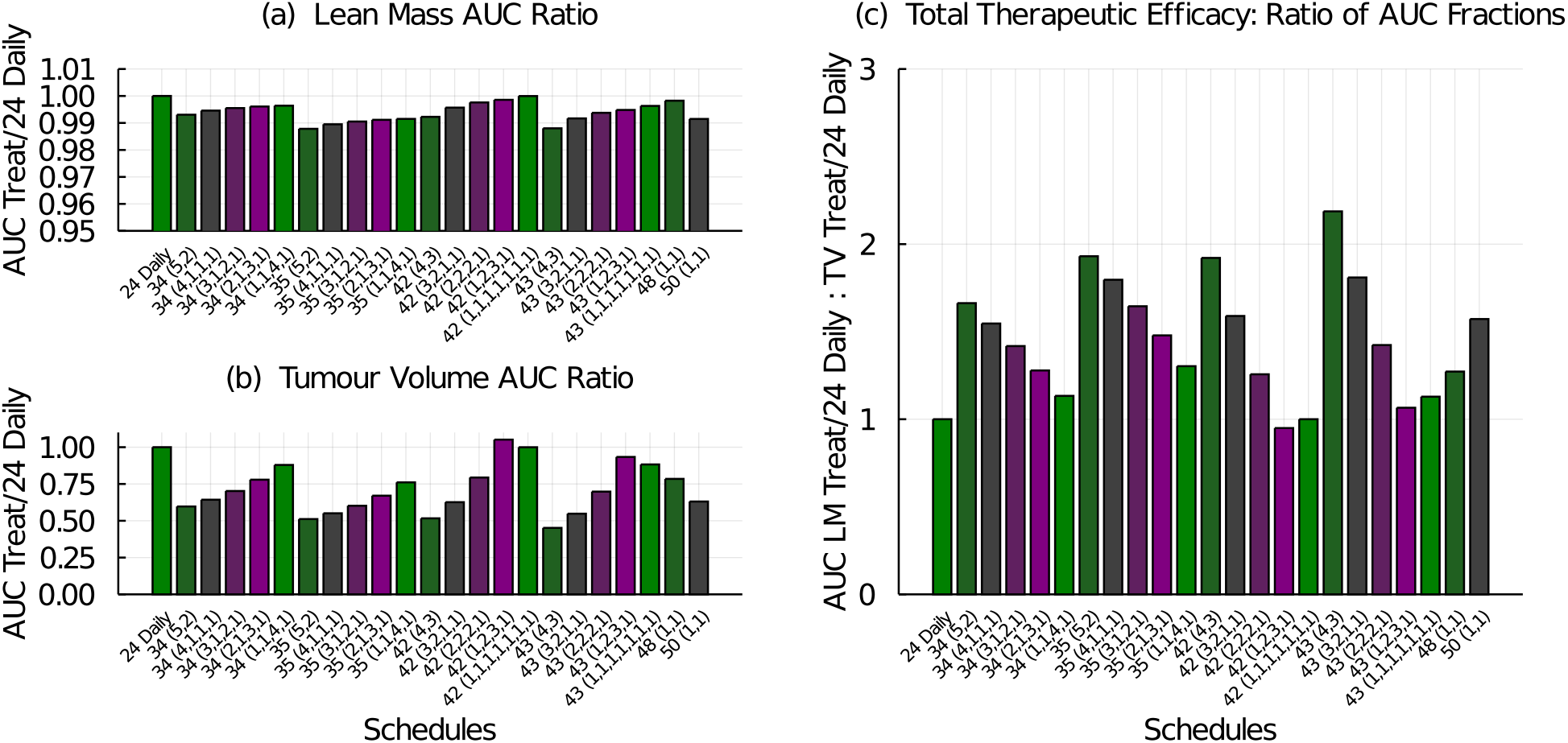
Treatment efficacy for the chemotherapy dosing schedules listed in Table 3. (a) The AUC ratio for lean mass compares the lean mass response from the tested schedule to that of the 24 mg/kg daily schedule. (b) The AUC ratio for tumour volume compares tumour volume response from the tested schedule to that of the 24 mg/kg daily schedule. (c) The total therapeutic efficacy is the ratio of the lean mass AUC ratio to the tumour volume AUC ratio.

Interestingly, delivering treatment every other day as in the 48 and 50 mg/kg (1-on, 1-off) schedules have improved tumour control over the 24 mg/kg daily schedule. The best total therapeutic efficacy schedule is again the 43 mg/kg (4-on, 3-off) schedule because it results in the best tumour control.

### Metronomic Schedules

We now consider schedules that deliver multiple doses per day to explore metronomic scheduling. Metronomic dosing attempts to maintain a more consistent level of the drug in the plasma in order to avoid the toxic side effects commonly seen in a maximum-tolerated-dose regime. Metronomic therapy is believed to target the endothelial cells and growing vasculature of a tumour whereas maximum-tolerated-dose therapy is believed to directly target the proliferating cancer cells [48].

The metronomic schedules examined include 1, 2, 3, 4, 5, or 6 doses equally spaced throughout each day, over a 4 week treatment period. The standard weekly dose of 168 mg/kg/week is broken up into equal doses and administered multiple times a day. For comparison, we use the 24 mg/kg daily schedule and the 35 mg/kg (5-on, 2-off) schedule.

The model predicted results of metronomic therapy are shown in Figure 11. Breaking the daily dose up over multiple doses per day results in a small improvement in lean mass preservation, Figure 11(a). It also results in less tumour control, Figure 11(b). Consequently, the multiple-doses-per-day metronomic schedules are predicted to be less therapeutically effective than the standard daily regime, Figure 11(c).

**Figure 11:**
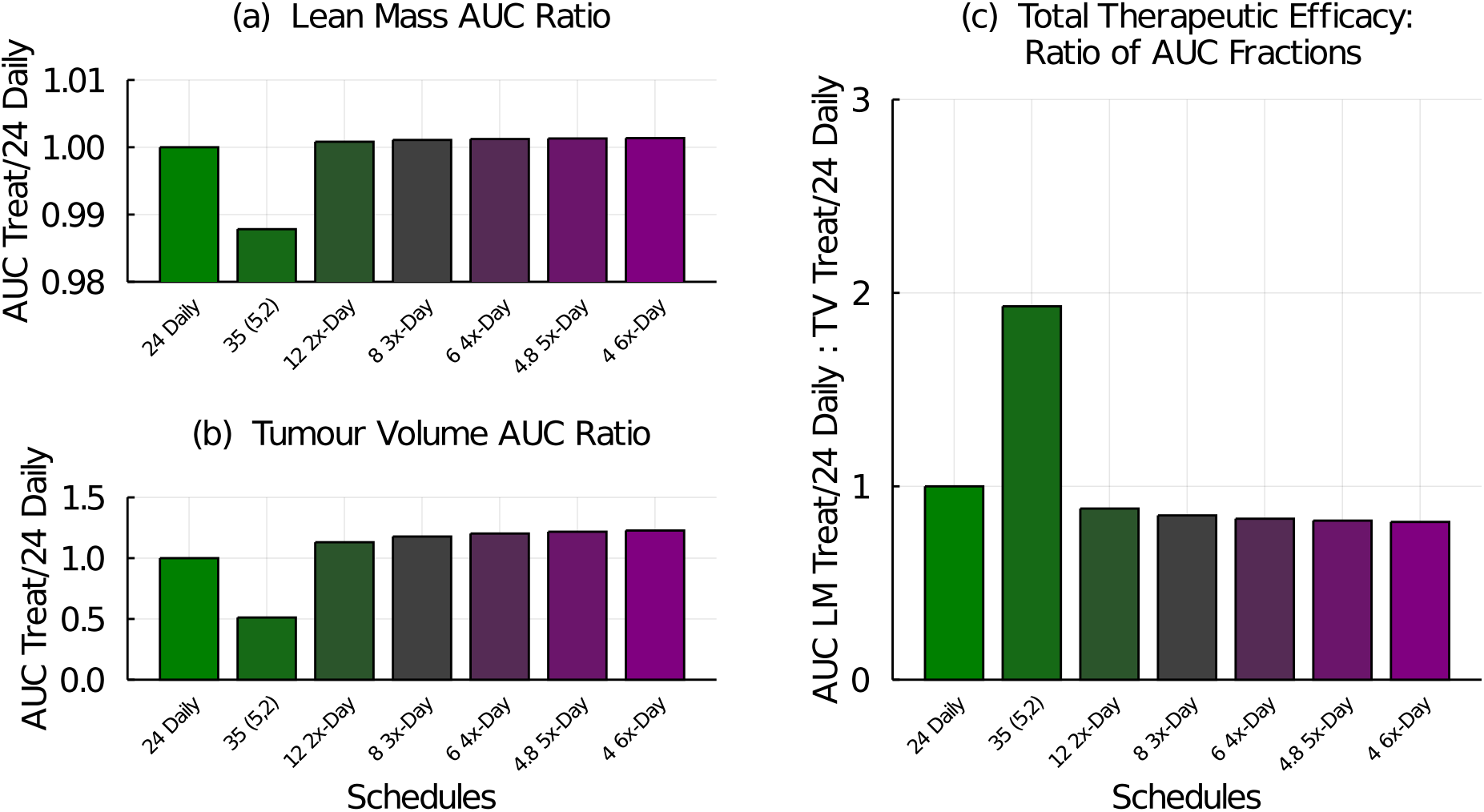
Treatment efficacy for metronomic chemotherapy schedules administering 168 mg/kg/week over 1, 2, 3, 4, 5, or 6 equal, and equally spaced, daily doses. Treatment efficacy is measured against the 24 mg/kg daily regime and the 35 mg/kg (5-on, 2-off) regime is included for reference. (a) The AUC ratio for lean mass compares the lean mass response from the tested schedule to that of the 24 mg/kg daily schedule. (b) The AUC ratio for tumour volume compares tumour volume response from the tested schedule to that of the 24 mg/kg daily schedule. (c) The total therapeutic efficacy is the ratio of the lean mass AUC ratio to the tumour volume AUC ratio.

### Cachexia in Aged Mice

Next we explore the effects of chemotherapy-induced cachexia in aged mice. The mice used in the experimental data and thus used to parameterize this model were 6 weeks old, and still in a phase of active growth. To explore the effects of aging on the chemotherapy-induced muscle loss, we simulate aged mice by starting our simulations with initial conditions matching a 2 year old. Further, we set the stem cell probability of self-renewing symmetric division feedback perturbation parameter, *p*_1_, to 98% of its nominal value which simulates an age-related decay in stem cell viability. This decrease instigates a gradual loss of muscle in the aged control of about 2.5%. The aged initial conditions are (*S*_0_, *M*_0_) = (275, 5333) mm^3^which gives a starting stem cell ratio of 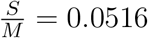 The 6-week old mouse simulations use initial conditions (*S*_0_, *M*_0_) = (268.5, 4732.5) mm^3^, giving a stem cell ratio of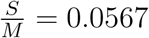.

To explore aging effects in chemotherapy response, we simulate several treatment regimes delivering 168 mg/kg/week of 5-FU for four weeks: 42 mg/kg (4-on, 3-off), 56 mg/kg (3-on, 4-off), 84 mg/kg (2-on, 5-off), 168 mg/kg (1-on, 6-off), 48 mg/kg every other day, and 8 mg/kg 3-times a day. The 24 mg/kg daily and 35 mg/kg (5-on, 2-off) schedules are included for comparison.

In Figure 12(a) the lean mass is simulated for the young (6 week) and old (2 year) controls. Whereas the young control is actively growing, the aged control experiences a small decline in lean mass. The effect of chemotherapy-induced cachexia transiently reduces the lean mass. Interestingly, the young simulations (solid lines) maintain the approximate mass observed at the start of treatment (about 10 g), but since the controls are still growing, this corresponds to a loss of about 1 g or 10% of lean mass. Conversely, the aged simulations (dashed lines) start at about 11 g and lose mass, due to the chemotherapy, until about 10% mass is lost. Both young and old recover after treatment cessation with the aged simulations lagging in recovery compared to the young. The stem cell ratio is transiently perturbed by treatment, Figure 12(b), with the aged hosts experiencing an downward drift in stem ratio due to the assumed decay in viability (*p*_1_).

**Figure 12:**
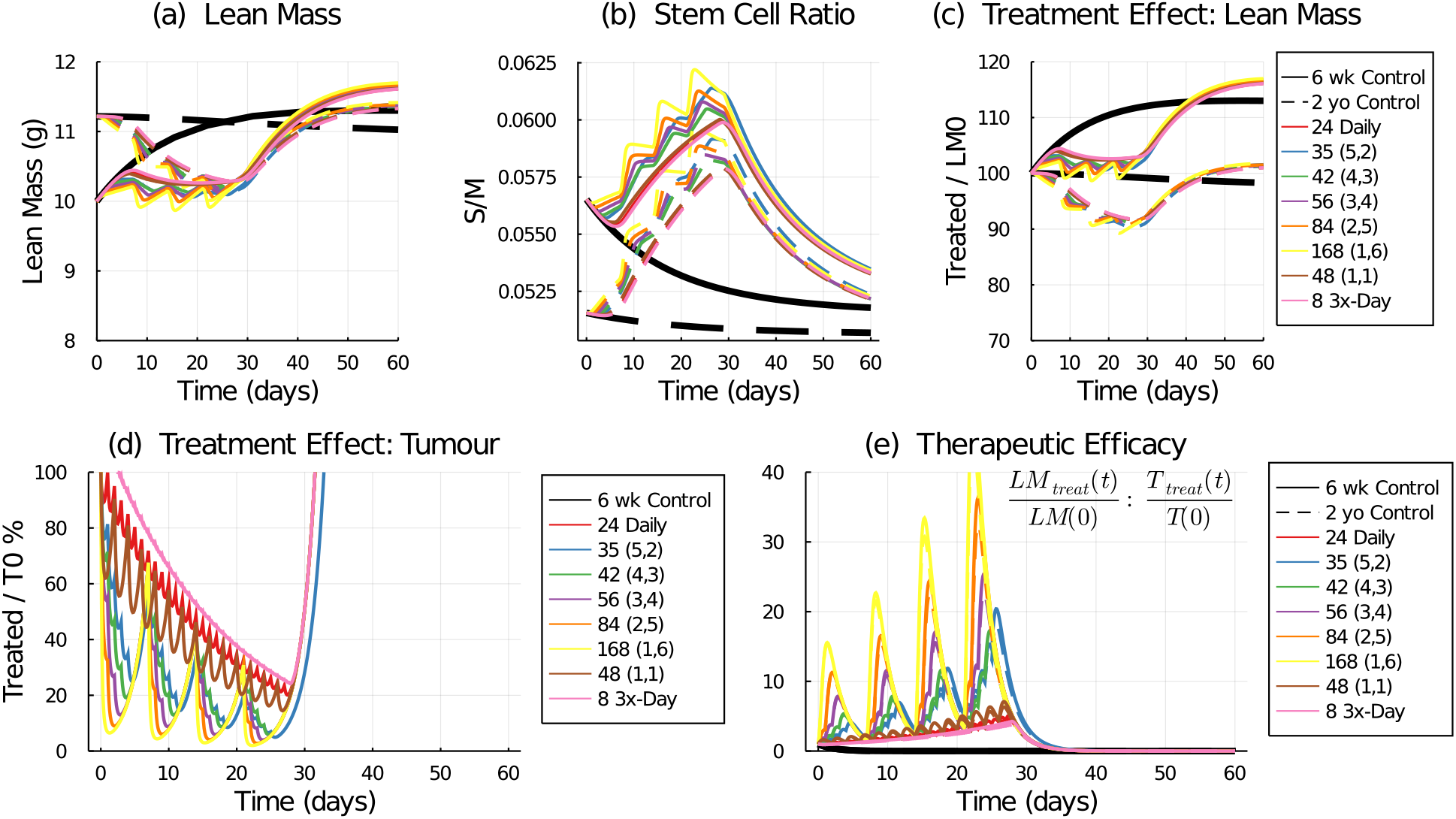
Comparison of chemotherapy effect on lean mass in a young (6 weeks old, solid lines) and old (2 years old, dashed lines) simulated host. Predicted lean mass response to treatment (a) and the resulting stem cell ratio (b). Treatment effect on lean mass (c) defined as the ratio of treated lean mass to initial lean mass. Treatment effect on tumour volume (d) defined as the ratio of treated tumour volume to initial tumour volume. Therapeutic efficacy (e) defined as the ratio of the treatment effect on lean mass to the treatment effect on tumour volume.

The treatment effect on lean mass, Figure 12(c), again shows about a 10% reduction in mass by the end of treatment compared to the young or old time-matched controls. Treatment effect on the tumour, Figure 12(d), is the same for young and old as no interaction between the host and the growing tumour is assumed in this model. The best treatment response is achieved by the maximum-tolerated-dose regime of 168 mg/kg delivered once a week. And finally, the therapeutic efficacy, Figure 12(e), shows the ratio of the treatment’s effect on lean mass to the treatment’s effect on tumour volume - as measured against their initial conditions. In these simulations, the young and old hosts are predicted to have similar responses and thus therapeutic efficacies to the tested schedules.

Figure 13 shows the AUC measures for lean mass and tumour volume response to treatment for the young and aged hosts, as measured against the 24 mg/kg daily treatment regime. The aged host is predicted to experience slightly less mass loss than the young host, Figure 13(a). The tumour response is independent of the host age, and the best treatment response is achieved by the maximum-tolerated-dose strategy of 168 mg/kg delivered weekly (1-on, 6-off), see Figure 13(b). Total therapeutic efficacy, Figure 13(c), again shows the best treatment responses are those that deliver the most drug in the shortest amount of time, following the maximum-tolerated-dose strategy. Delivering drug daily, every other day, or three times a day, all perform comparably. No significant age-related differences are captured by the AUC measures which integrate the treatment responses over the 28 days of the schedule.

**Figure 13:**
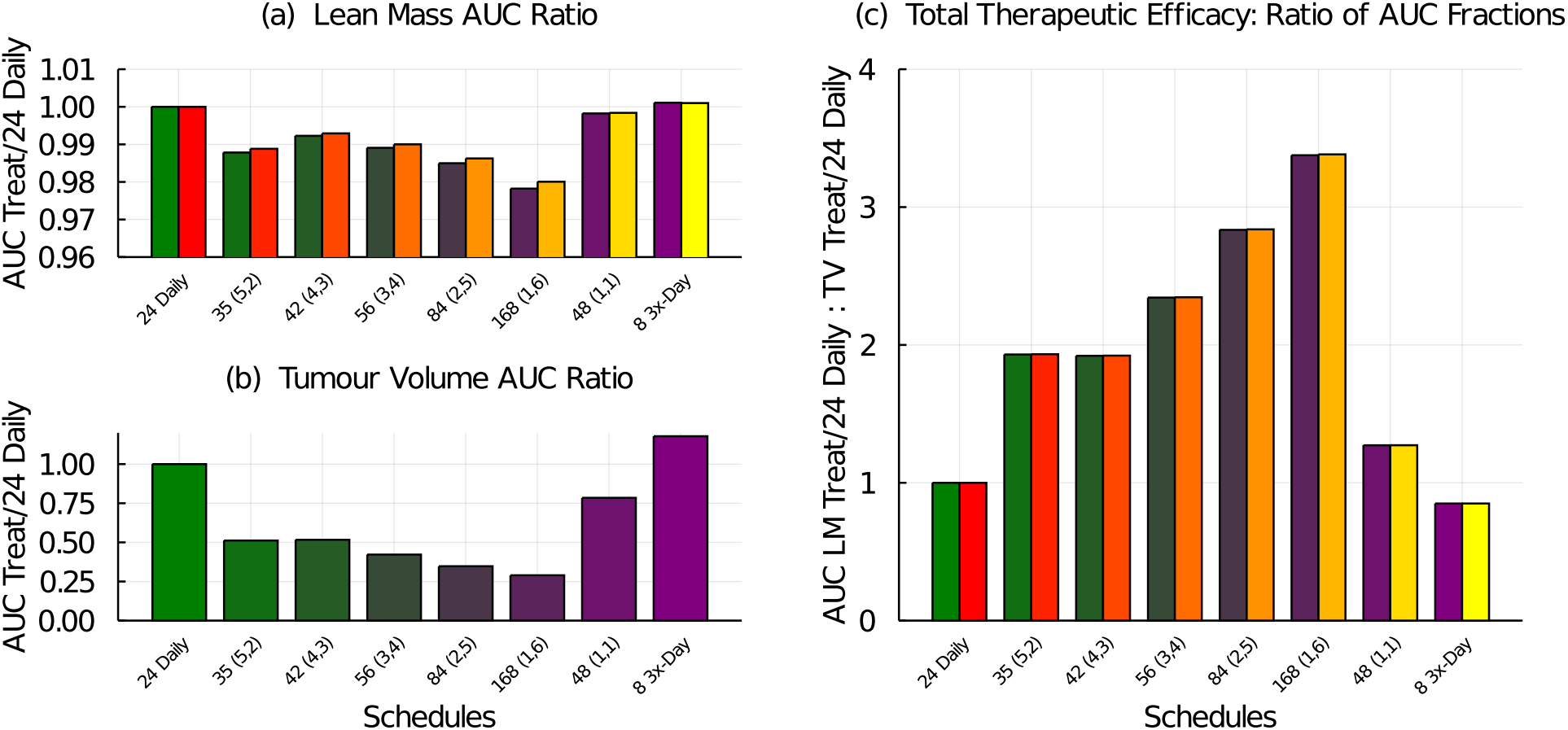
Treatment efficacy for young (6 weeks, green-purple bars) and old (2 years, red-orange bars) simulated hosts exposed to chemotherapy schedules administering 168 mg/kg/week in 1, 2, 3, or 4 doses per week, as well as a 3-times a day schedule and an every-other-day schedule. Treatment efficacy is measured against the 24 mg/kg daily regime and the 35 mg/kg (5-on, 2-off) regime is included for reference. (a) The AUC ratio for lean mass compares the lean mass response from the tested schedule to that of the 24 mg/kg daily schedule. (b) The AUC ratio for tumour volume compares tumour volume response from the tested schedule to that of the 24 mg/kg daily schedule. (c) The total therapeutic efficacy is the ratio of the lean mass AUC ratio to the tumour volume AUC ratio.

### Morphine and Chemotherapy-Induced Cachexia

Concomitant use of morphine with 5-FU has been shown to decrease the clearance and elimination rates of the chemotherapy agent in mice [49]. Plasma levels were elevated after morphine administration during 5-FU treatment. In particular, the clearance rate of an intravenus bolus of 100 mg/kg 5-FU was reduced from 54 to 28 ml/min/kg (a reduction of about 52%) and the elimination half-life was increased from 6.9 to 12.2 min (an increase of about 177%) by prior morphine administration [49]. Continuous infusion of 5-FU after morphine administration was also found to have a 35% reduction in clearance rate. Morphine tolerance was shown to reduce the effects on 5-FU clearance, as after prolonged exposure, higher doses of the opiate were needed to obtain the same level of pain management and to raise the 5-FU plasma levels. Thus it is possible that in treating pain, off-target chemotherapy effects on muscle may be inadvertently exacerbated, ultimately leading to further increased pain levels and decreased quality of life.

We explore the effects of morphine on our chemotherapy-induced cachexia model by reducing the 5-FU pharmacokinetic transit and clearance parameters (*k*_21_, *k*_12_, and *k*_10_) by 35%. We assume this effect is permanent, and ignore any acclimatization that may occur with prolonged morphine use. The reduction in transit and clearance rates increase the levels of 5-FU in the model compartments, and thus increase the *τ* -day average exposure function, see Figure 14.

**Figure 14:**
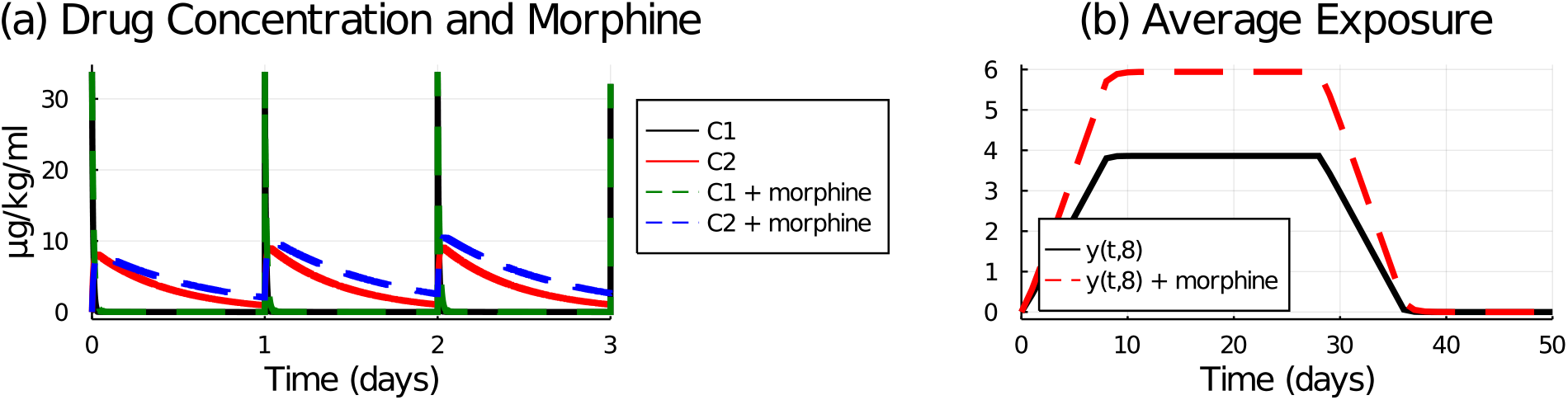
Drug concentration in plasma and tissue model compartments (a) and the average exposure function with *τ* = 8 (b). Solid lines are control (no morphine) and dashed lines are with morphine where model parameters *k*_21_, *k*_12_, and *k*_10_ were reduced by 35%.

Model simulated lean mass and tumour volume under 5-FU treatments and adjuvant morphine administration are shown in Figure 15. Compared with the control (no morphine), the morphine addition is predicted to cause significant and dangerous lean mass loss for all tested schedules, Figure 15[a]. It also improves tumour control, Figure 15[b]. And as expected, the schedule that preserves the most lean mass is the metronomic (3 times a day) schedule, Figure 15[c], while the best tumour control and thus highest total therapeutic efficacy is achieved with the maximum-tolerated-dose (168 mg/kg delivered 1 day a week) schedule, Figure 15[d,e].

**Figure 15:**
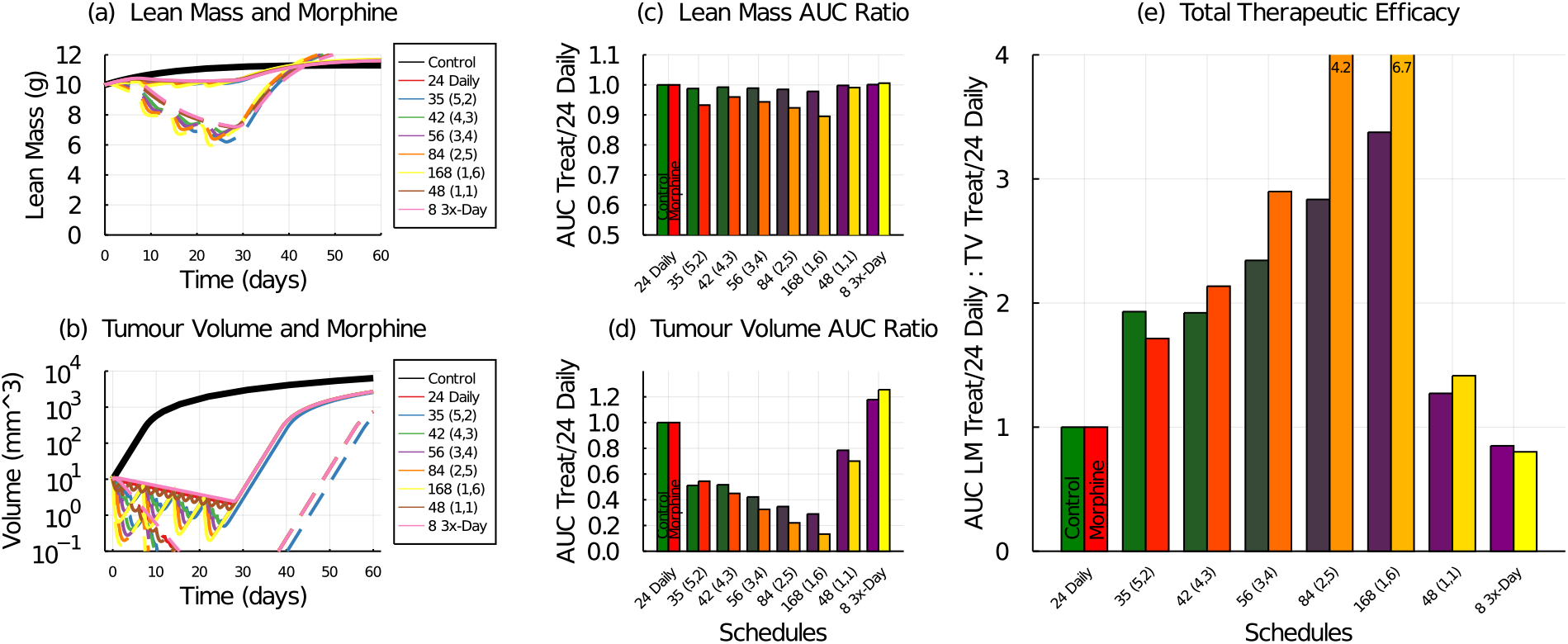
Model simulation of various 5-FU dosing schedules with and without adjuvant morphine administration. Lean mass (a) and tumour volume (b) response to various 5-FU schedules with (dashed lines) and without (solid lines) morphine. Treatment efficacy without (green-purple bars) and with (red-orange bars) morphine for chemotherapy schedules administering 168 mg/kg/week in 1, 2, 3, or 4 doses per week, as well as a 3-times a day schedule, and an every-other-day schedule. Treatment efficacy is measured against the 24 mg/kg daily regime, and the 35 mg/kg (5-on, 2-off) regime is included for reference. (a) The AUC ratio for lean mass compares the lean mass response from the tested schedule to that of the 24 mg/kg daily schedule. (b) The AUC ratio for tumour volume compares tumour volume response from the tested schedule to that of the 24 mg/kg daily schedule. (c) The total therapeutic efficacy is the ratio of the lean mass AUC ratio to the tumour volume AUC ratio.

These model simulations represent a worst-case scenario, where morphine usage is continuous and increasing to account for any acclimatization effects. The model predictions, however, demonstrate that inadvertent interference between pain-management drugs and chemotherapy regimes may exacerbate both on-target and off-target effects of the drug, and thus warrant further investigation.

## 5 Discussion

Since muscle loss is correlated with poor survival rates, understanding the mechanisms of cachexia, including chemotherapy-induced cachexia, and designing treatments that aim to preserve muscle mass in addition to tumour control, are of great importance. This work presents a first attempt to mathematically model chemotherapy-induced muscle loss in a manner that captures the nonlinear dose-dependence observed in experimental data [37]. The model consists of a two-compartment pharmacokinetic system for 5-FU, a two-compartment stem-cell lineage system for muscle mass, and a tumour growth equation, all coupled together. The growth equations are assumed to be negatively affected by the chemotherapy concentration in a secondary (muscle tissue or tumour mass) compartment. The stem cell and muscle compartments in particular, suffer from reduced successful proliferation via a cubic function of a *τ* -day average drug exposure which captures the initial delay in mass loss observed in the dataset.

The parameters that couple our model together, *τ* and *R*_*d*_, were estimated by fitting simulations of all doses to the data for either the daily or 5-day dosing schedules. The best-fit parameters for all schedules were then chosen as the average of the best fits for the two experimental dosing schedules. A dynamic sensitivity analysis was performed as the chemotherapy effect is transitory. The sensitivity results for *R*_*d*_ highlight how the drug holidays in the (5-on, 2-off) schedule significantly reduce treatment effects on lean mass.

With this model we explore various dosing schedules and compare their effectiveness by examining area-under-the-curve based metrics for lean mass and tumour volume response to treatment, and for the total therapeutic efficacy. In general, lean mass is best preserved by following a metronomic schedule (doses once or more a day) and tumour control is best achieved by following a maximum tolerated dose schedule (doses once a week).

Lastly, we used our model to explore potential confounding effects of aging and morphine usage. By the end of our simulated treatment, the old and young mice lost approximately the same amount of lean mass, and experience similar disruptions to their stem cell ratios. Simulated morphine usage drastically increased the effects of 5-FU, causing a much more significant lean mass loss along with a much improved tumour control.

Together, this work highlights the difficulty in defining an optimal treatment schedule in terms of reducing tumour volume while also maintaining lean muscle mass. Across all simulations, the maximum tolerated dose schedule performed best for tumour control and worst for lean mass preservation. The metronomic schedules (daily or more frequent) performed best for lean mass preservation but worst for tumour control. Thus, defining an optimal schedule consists of a weighted balance between improved tumour control and preserved lean mass.

In mathematical modelling of chemotherapy it is standard to assume, as we do here, that cell kill is proportional to the drug concentration. This necessarily means that larger doses will affect larger cell death. But this type of model cannot explain the reported successes of metronomic therapy with their improved toxicities, lower costs, and ease of use [50]. The mechanisms of action of metronomic chemotherapy are thought to include inhibited angiogenesis, modulation of the immune response, and direct affects on tumour cells and stromal cells. To account for these mechanisms, a systemic (whole-body) modelling approach is required. Metronomic chemotherapy is promising for personalized treatment plans, especially in the elderly or those suffering from cachexia, who may not complete the full course of a maximum tolerated dose regime.

The modelling and simulation work presented here is a first attempt to capture the non-linear dose-response of skeletal muscle to cancer chemotherapy treatment. One simplification made was to neglect immune cells, which play significant roles in advancing cachexia by promoting inflammation, and in maintaining muscle homeostasis by regulating cell turnover and coordinating repair and remodelling [51]. Such immune cells are sensitive to chemotherapy treatment and their loss can exacerbate muscle loss and dysfunction. Consideration of these important aspects are proposed for future study.

Lean mass body composition can be negatively affected by aging or cancer cachexia, and is related to chemotherapy metabolization and toxicity [52]. Patient-to-patient variations in muscle mass are not accounted for in treatment planning and may contribute to treatment outcome variability [53]. Body composition is thus a potential factor to integrate into the design of patient-specific treatment plans.

To the authors knowledge, there are currently no approved therapies for cancer cachexia or chemotherapy-induced cachexia even though muscle loss is known to be a significant factor in treatment toxicity and patient quality of life. One potential treatment target to reduce the side-effects of chemotherapy is the gut microbiome. Adjuvant therapy designed to target the microbiota and dampen the inflammatory cascade instigated by cancer and cancer treatment has been proposed [54].

Taken together, the immune response, body composition, and microbiome can inform the whole-body health level of a patient, which can be integrated into patient-specific treatment planning, including metronomic schedules. These exciting prospects highlight the systemic nature of cancer and the multi-faced dysregulation that occurs from both the disease and its treatment. Additionally, they highlight a need for more systemic modelling at the host level that incorporate such factors as inflammation and the immune response (not just tumour cytotoxicity), muscle mass, and gut microbiome, as we move into personalization of treatment planning.

## A Appendix Julia Code to Simulate Mathematical Model

Below is sample code to numerically simulate the described mathematical model with the *τ* -day average exposure function and all referenced model parameters.

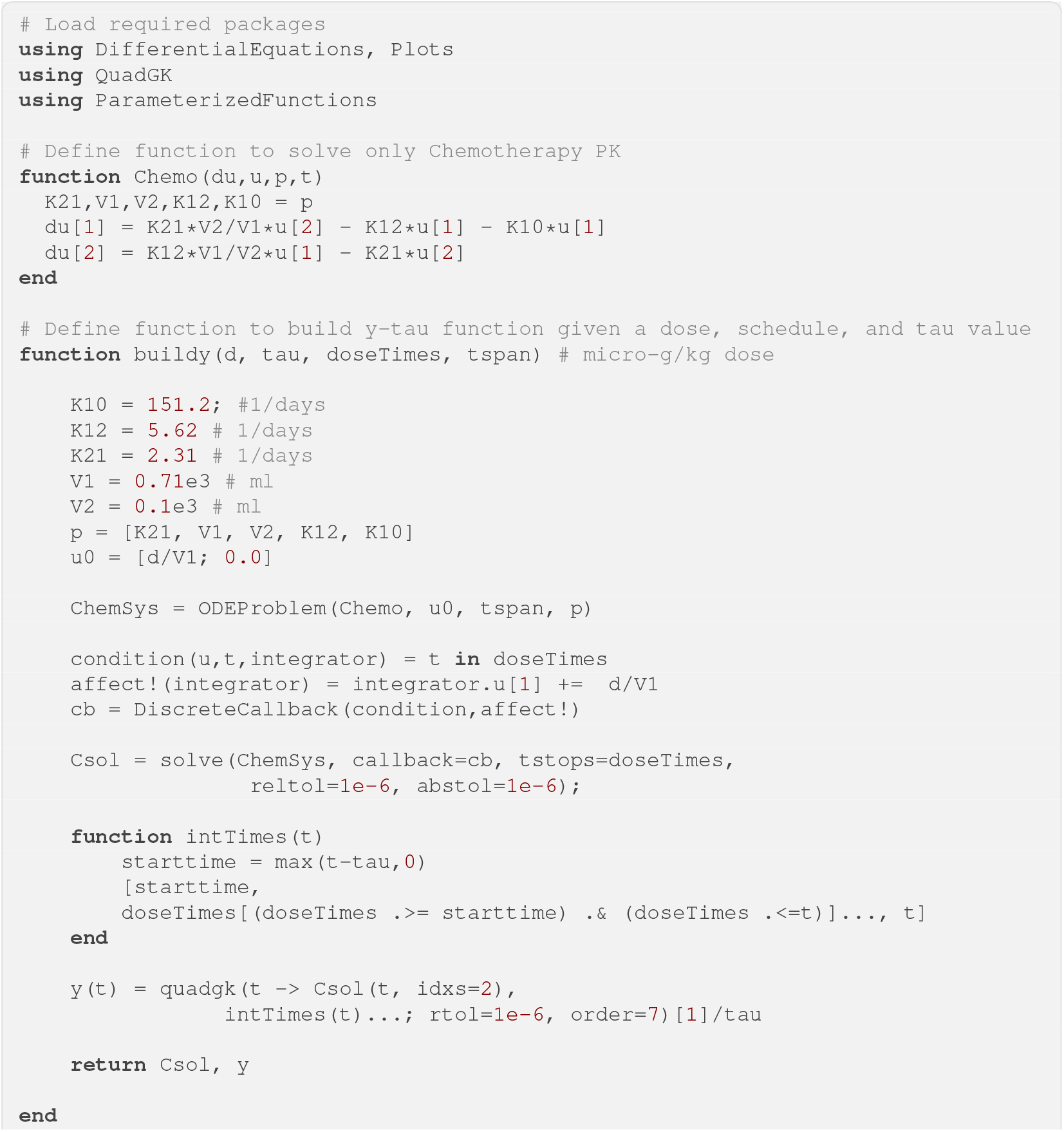
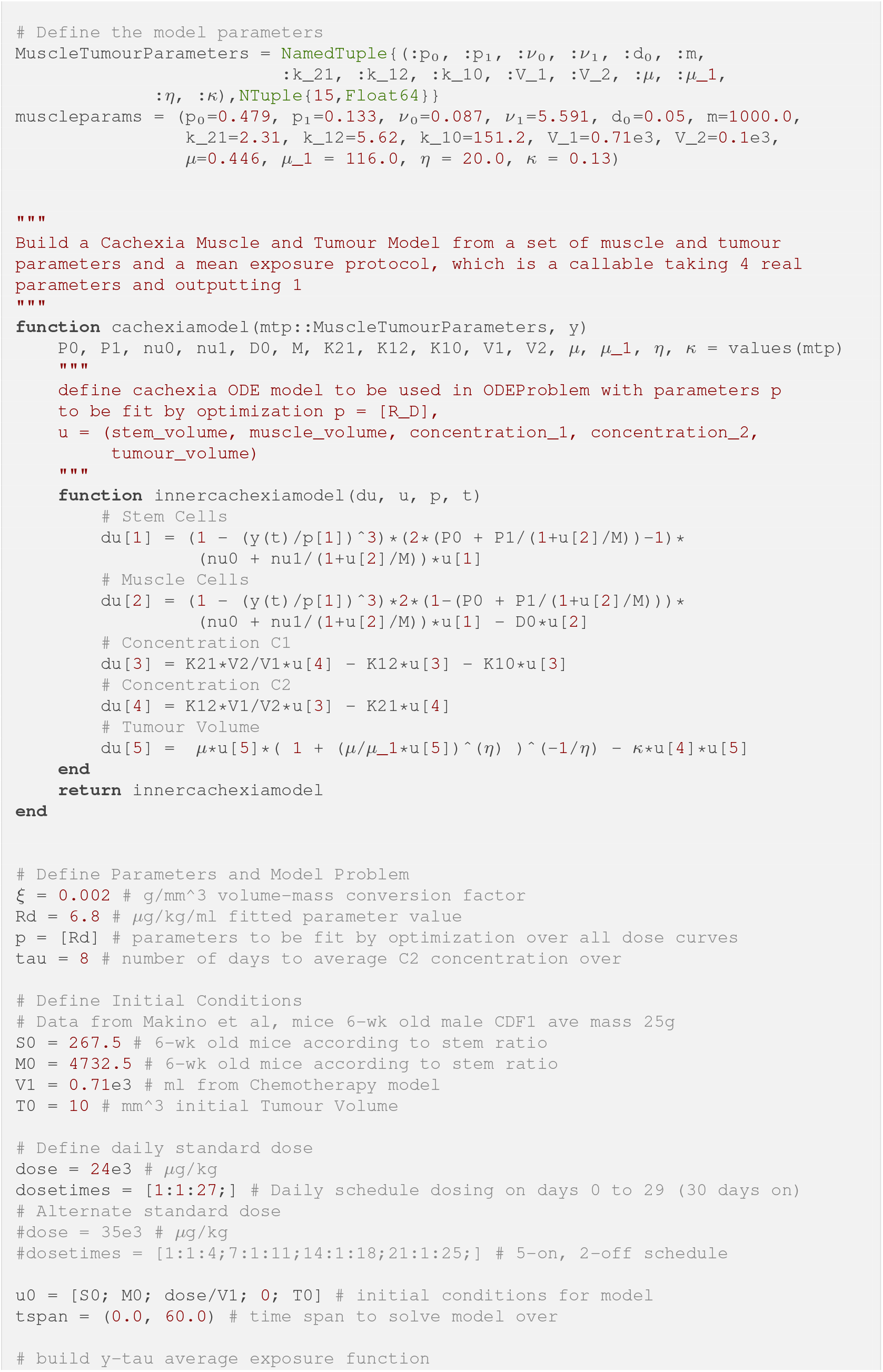
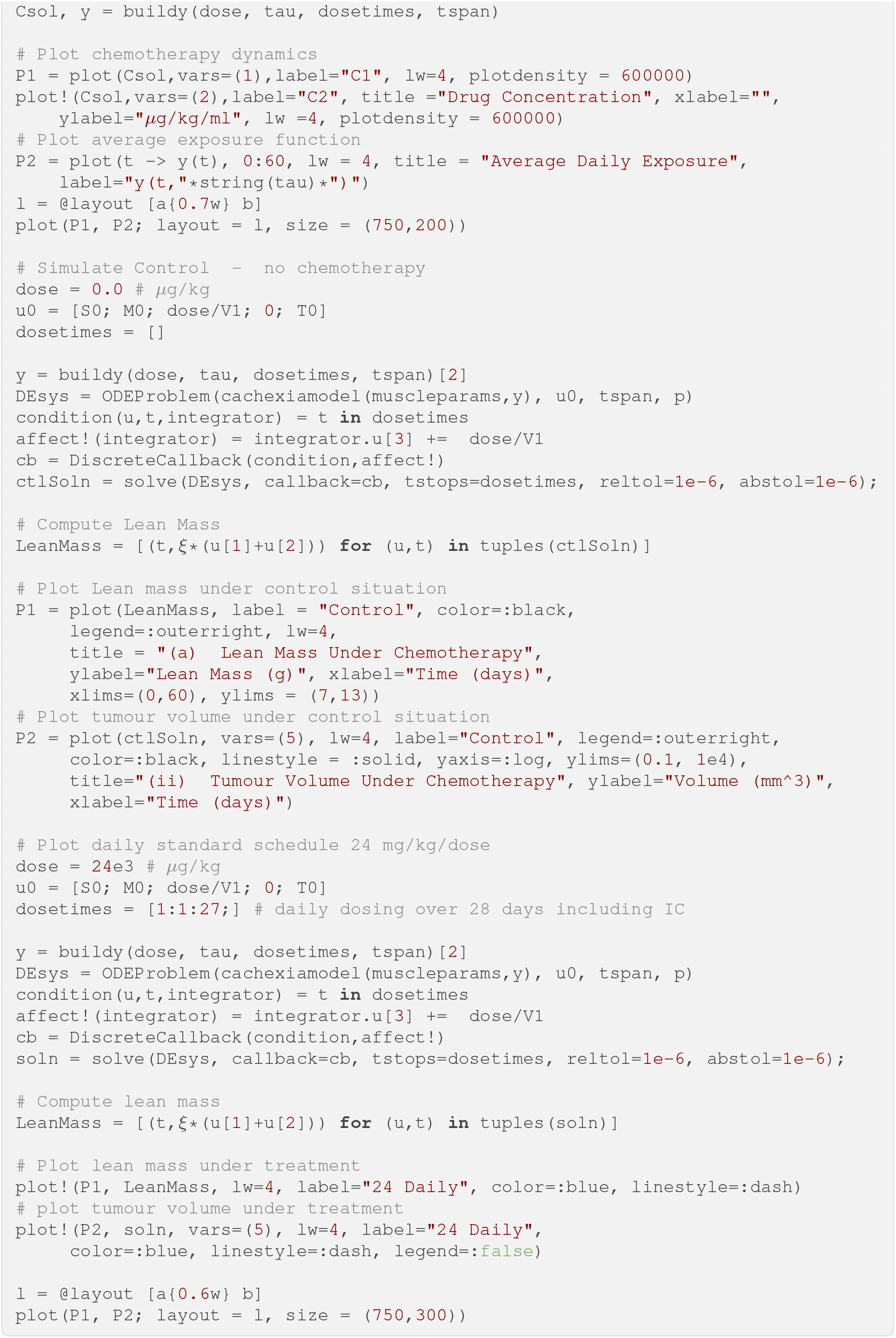

